# Cortical changes during the learning of sequences of simultaneous finger presses

**DOI:** 10.1101/2023.03.12.532251

**Authors:** Benjamín Garzón, Gunther Helms, Hampus Olsson, Claudio Brozzoli, Fredrik Ullén, Jörn Diedrichsen, Martin Lövdén

## Abstract

The cortical alterations underpinning the acquisition of motor skills remain debated. In this longitudinal study in younger adults, we acquired performance and neuroimaging (7T MRI) measures weekly over the course of 6 weeks to investigate neural changes associated with learning sequences of simultaneous finger presses executed with the non-dominant hand. Both the intervention group (*n* = 33) and the control group (*n* = 30) showed general performance improvements, but performance improved more and became more consistent for sequences that were intensively trained by the intervention group, relative to those that were not. Brain activity for trained sequences decreased compared with untrained sequences in the bilateral parietal and premotor cortices. No training-related changes in the primary sensorimotor areas were detected. The similarity of activation patterns between trained and untrained sequences decreased in secondary, but not primary, sensorimotor areas, while the similarity of the activation patterns between different trained sequences did not show reliable changes. Neither the variability of activation patterns across trials, nor the estimates of brain structure displayed practice-related changes that reached statistical significance. Overall, the main correlate of learning configural sequences was a reduction in brain activity in secondary motor areas.

## Introduction

The ability to move is indispensable for key survival functions, so much so that the brain devotes vast resources to generating movement. Beyond survival, professional achievement and social recognition are often reliant on the acquisition and subsequent production of sophisticated motor behaviors (for instance, in the case of craftspeople, athletes, or musicians). Even mundane activities such as typing on a computer require complex motor programs. For these reasons, understanding the process of motor skill acquisition has been a long-standing topic in psychology and cognitive neuroscience. Nonetheless, many outstanding questions remain, and a key one is how the observed behavioral changes are related to the alterations known to occur at multiple neurophysiological levels (Zatorre et al. 2012; Sampaio-Baptista et al. 2013; Fields 2015; Jensen and Yong 2016; Makino et al. 2016; Krakauer et al. 2019).

There is ample evidence from functional Magnetic Resonance Imaging (fMRI) studies of learning-related changes in activation in primary and secondary motor cortices as well as in subcortical areas (Grafton et al. 1995; Karni et al. 1995, 1998; Penhune and Doyon 2002; Floyer-Lea 2005; Lehéricy et al. 2005; Dayan and Cohen 2011). However, there are discrepancies among studies regarding whether activity in these areas increases, decreases, or shows more complex non-monotonic patterns of change with practice. It also remains unclear whether the changes occur in the primary sensorimotor cortices or exclusively in secondary areas (Xiong et al. 2009; Ma et al. 2010; Huang et al. 2013; Wiestler and Diedrichsen 2013; Berlot et al. 2020). Recent work made significant progress in resolving these disagreements with a preregistered long-term longitudinal study of subjects practicing a finger-sequence production task (Berlot et al. 2020). This study found decreases in activation during performance relative to rest in trained compared with untrained sequences in the dorsal premotor cortex and the anterior superior parietal lobule. Activation in primary sensorimotor cortex remained constant. Berlot and colleagues (2020) also reported changes of the multivariate activation patterns for the execution of trained sequences in secondary, but not primary, regions (Huang et al. 2013; Wiestler and Diedrichsen 2013; Berlot et al. 2020).

Learning classic finger-sequence tasks primarily requires assembling elements from the repertoire of previously learned actions instead of creating novel continuous movements (Krakauer et al. 2019; Wong and Krakauer 2019). According to this interpretation, what is learned in this type of task is to select rapidly the appropriate, already-learned, discrete actions in the correct order. Learning may thus primarily occur in movement planning and action selection – an interpretation consistent with observations of learning-relate changes in secondary but not primary sensorimotor cortices (Yokoi 2019). This perspective is compatible with a hierarchical architecture of the representations underlying motor skill, with associative areas encoding chunks and sequences of elementary motor components or particular component features like timing or spatial organization (Diedrichsen and Kornysheva 2015). Nevertheless, it remains unknown whether learning sequences of more difficult discrete movements (i.e., movements that are initially challenging because they have not been practiced previously) is associated with the same pattern of activation over time (i.e., stability in the primary sensorimotor cortices and reductions in the associative cortices). That is, the primary sensorimotor cortices may also be involved in learning such sequences of movements. Here we addressed this issue by developing a configural sequence task, akin to playing short sequences of piano chords, which naïve subjects had to learn to execute with their non-dominant (left) hand (Figure 1).

**Figure 1.**
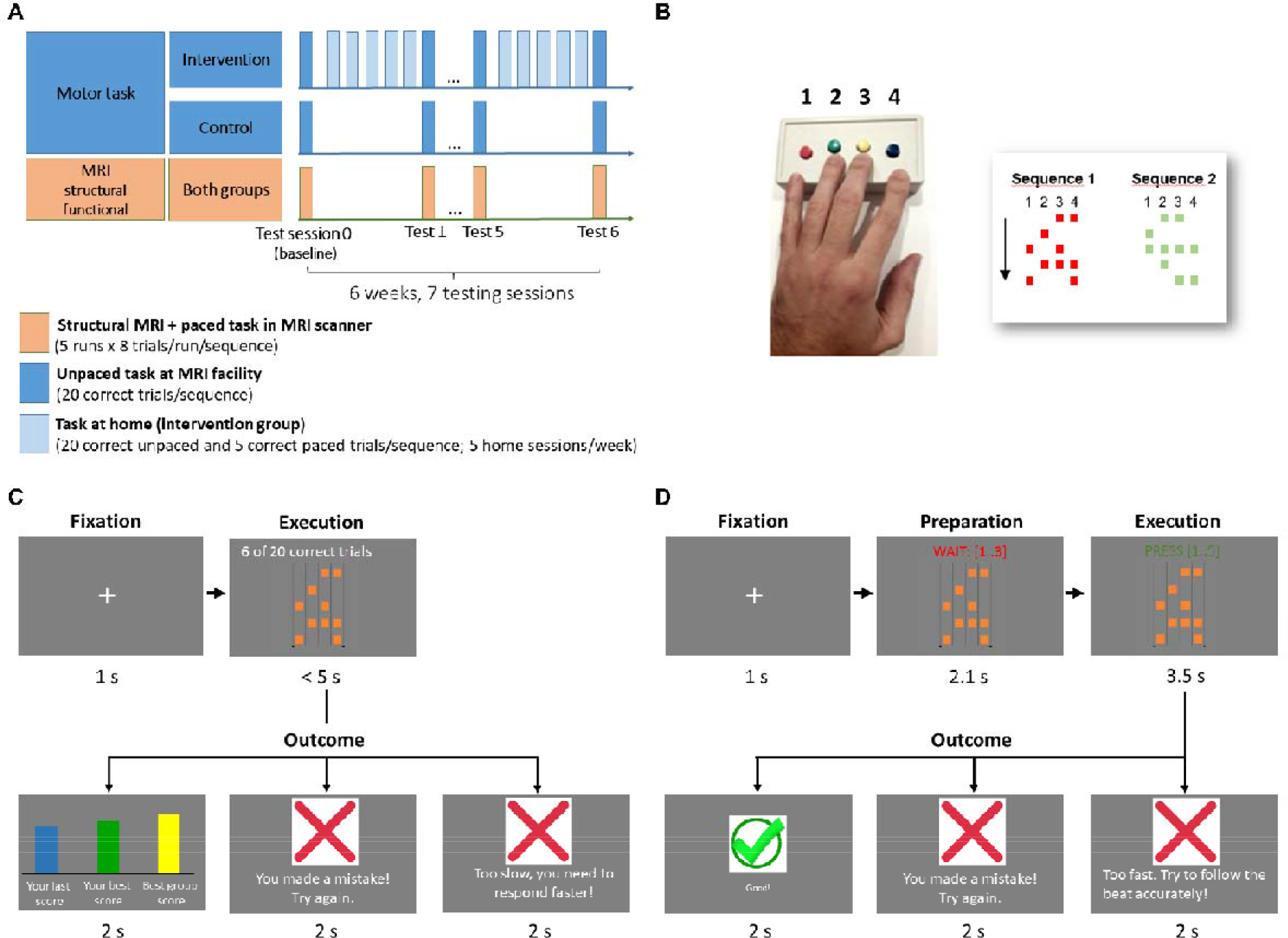
Design and task. A) The longitudinal study spanned a 6-week period, with 7 testing sessions. Participants were tested behaviorally once a week at the MR facility, both outside the scanner (unpaced task) and inside the scanner (paced task) while undergoing functional MRI. Structural images were also acquired. In between testing sessions, subjects in the intervention group practiced at home, 5 times a week. B) Participants practiced and were tested on a discrete configural-response sequence task, in which they had to execute different sequences of finger movements with 5 combinations (chords) of up to 4 fingers with their left (non-dominant) hand upon seeing a cue. Subjects used a button box and each finger except the thumb had one button assigned to it. Each button that had to be pressed was depicted as a square, and button combinations were arranged in 5 rows, to be executed from top to bottom. Subjects practiced two versions of the task, both starting with a short fixation, after which a cue was shown depicting the sequence to be executed. In the unpaced version of the task (C), subjects were asked to execute the sequence correctly and as fast as possible. They received feedback regarding correctness and execution speed, relative to their past trials and to the best score in their (fictitious) peer group. A sequence had to be repeated until 20 correct trials were achieved before moving on to the next. In the paced version (D), a counter indicated a beat (0.7 s) that participants had to follow when pressing the chords. Participants received feedback regarding whether the sequence was correct and in sync with the beat, and they had to achieve 5 correct trials to move on to the next sequence. In the scanner, participants were tested on the paced version of the task but no feedback was provided, and a different sequence was presented on each trial, with two consecutive trials never presenting the same sequence. See Methods for details about the design and the task.

Beyond functional changes, human neuroimaging has also revealed learning-related alterations in brain structure. Estimates of gray and white matter structure obtained with MRI display differences between adult human experts and non-experts in brain regions that are relevant to their domain of expertise (Amunts et al. 1997; Maguire et al. 2000; Gaser and Schlaug 2003; Bengtsson et al. 2005; de Manzano and Ullén 2018). In longitudinal designs, changes in such structural properties can be detected following weeks or months of practice to acquire skills such as juggling or speaking a new language (Draganski et al. 2004, 2006; Scholz et al. 2009; Mårtensson et al. 2012; Lövdén et al. 2013; de Lange et al. 2017). Some evidence indicates that such structural changes in gray matter may be non-monotonic, with initial increases followed by partial normalization during motor learning (Wenger et al. 2017).

Several researchers have attempted to integrate these findings of learning-related structural changes with the functional changes, and with related observations in animal models, like cortical map reorganization (Molina-Luna et al. 2008; Reed et al. 2011) or the formation and stabilization of selected synapses (Xu et al. 2009; Yang et al. 2009), under the umbrella of an exploration-selection-refinement (ESR) theory (Kilgard 2012; Makino et al. 2016; Lindenberger and Lövdén 2019; Lövdén et al. 2020). This conceptual model draws from early ideas of variation and selection within neural populations (Edelman 1987; Changeux 1989) and predicates that remodeling of circuits in motor skill learning follows three phases. Initially, circuits that may elicit potentially adequate movements are randomly recruited (exploration), leading to a high activation extent that prompts structural changes in those circuits. Subsequently, the neural ensembles supporting movements that are reinforced persist, while superfluous circuitry is pruned (selection). Finally, the selected circuits are fine-tuned as the optimal behavior is repeated, resulting in stable long-term memories and a slow development of precision and consistency of performance (refinement).

The ESR theory generates hypotheses that are testable in humans with MRI: (1) The variability of neural representations (i.e., activity patterns) across trials corresponding to the same intended action should be initially high during exploration and then decrease rapidly after selection; (2) the overall neural activity level should be initially high and then decrease during learning; and (3) the ESR process should give rise to growth of regional structure in brain regions controlling the learned movement during the exploration phase (expansion), followed by a partial retraction (renormalization) after the best circuit for the task has been selected (Lindenberger and Lövdén 2019). In the present study we sought to test these preregistered predictions (https://osf.io/48meb; https://osf.io/x4c9b). Under the assumption that the task that we developed demands learning of both novel elementary movements (simultaneous multiple finger presses) and sequential combinations that have not been a prominent part of the subjects’ behavioral repertoire before, we predicted these results in both primary and secondary sensorimotor areas. In analyses that were not preregistered, we also probed past findings of reductions in behavioral variability across trials of the same intended action and examined how training affects the similarity between activation patterns elicited by different movement sequences.

To test these hypotheses, we randomly allocated 70 healthy right-handed younger adults to either an intervention group, which trained the task at home 5 times a week during a period of 6 weeks, or to a control group (Figure 1). Once a week, both groups were scanned with structural and functional MRI, and tested inside and outside of the scanner on trained and untrained sequences. With this design, we could probe sequence-specific learning by comparing changes in performance and brain activity between trained and untrained sequences, controlling for the effects of repeated testing. Furthermore, comparison between the intervention and control groups enables assessing transfer of training effects to untrained sequences (i.e., sequence-general effects of training) and to investigate the effects of learning on brain structure. We measured baseline brain structure and activity before any substantial pretraining. Some discrepancies between previous studies could stem from whether or not they included pretraining, which could be associated with some early changes before the baseline scans. Training of the sequences took place with both unpaced instructions (i.e., subjects were required to complete the sequences as fast as possible) and paced instructions (i.e., subjects were required to execute the discrete movements following a predefined tempo). The paced condition was administered during fMRI to rule out that potential training-related activity changes were driven simply by changes in motor output.

## Methods

### Participants and recruitment procedures

Subjects were recruited via advertisements on a recruitment website and in an online newspaper, and with flyers in the local area. Volunteers who appeared to fulfill study criteria upon a screening via phone were invited to attend an introduction meeting, in which a researcher provided information concerning the study. Subjects who agreed to partake in the study signed a consent form. They were then asked to complete questionnaires focused on study criteria, the Edinburgh Handedness Inventory, and an 18-item version of Raven’s Progressive Matrices test. Next, subjects tried a short demo of the practice routine (with simpler sequences and fewer trials than they would encounter in the actual experiment) so that they could ask questions and familiarize themselves with the task and the button box. The researcher ensured that participants understood the task. Participants who fulfilled all study criteria (see Supplementary Table 1) received an invitation to take part in the study.

The recruited subjects (*n* = 70; age = 20-30 years, right-handed, MRI eligible, no previous experience of fine-motor skill acquisition involving the left hand) were randomly allocated to either an intervention (*n* = 35) or a control group (*n* = 35), matched for sex and score in the Raven’s matrices test. Participants were informed that they would be randomly allocated to two different groups, but their group membership was only disclosed after finishing the baseline session. To be able to fit all the planned non-linear trends over test sessions, we excluded participants completing less than 4 scanning sessions for the analyses reported here, leaving 33 intervention subjects and 30 controls. The two groups considered in the final analyses did neither differ statistically significantly in sex (M/F intervention group: 12/21; control group: 12/18; Kruskal-Wallis χ^2^(1) = 0.09, p = 0.78) nor in performance on the Raven’s Progressive Matrices test (intervention: mean = 10.7, SD = 2.9; control: mean = 10.8, SD = 3.2; two-tailed t(59.14) = 0.13, p = 0.90). Supplementary Figure 7 shows a flow chart for the recruitment, attrition, and exclusions in the study.

Subjects received financial compensation for each training session and for each MR exam, and a scheme of rewards and penalties was implemented to incentivize compliance and effort. Participants in the control group were paid up to 5850 Swedish crowns (SEK) if they completed all the sessions, whereas participants in the intervention group were paid up to 8350 SEK due to the additional dedication required by the home training sessions.

### Ethics statement

The study was reviewed and approved by the Ethical Review Board in Stockholm (Case number: 2018/1620-31/2). All the participants gave their informed consent.

### Motor task

The task was a configural sequence learning task, which requires the execution of a sequence of key combinations that need to be pressed simultaneously with fingers of the non-dominant hand, as when playing a sequence of chords on a piano keyboard. The sequences had 5 combinations of between 1 and 4 fingers, excluding the thumb, and subjects used a button-box with 4 buttons (Current Designs, Philadelphia, PA, www.curdes.com) to execute them (Figure 1 B). Depending on the type of session, subjects went through between 3 and 6 different sequences. The sequences were associated with different colors to make it easier for the participants to identify them and recognize when there was a sequence change during the session. At the beginning of each session, the software showed instructions on the screen. The subject triggered the start of the task by pressing the space button. Each trial started with a fixation cross on screen (1s). A representation of a sequence was then displayed (5 rows with squares denoting which fingers to press in the following order from left to right: pinky, ring, middle, index) that the participant had to execute. The task was implemented with the Psychopy library (psychopy.org) for python (python.org).

After the baseline session, subjects were told which group they had been allocated to and intervention subjects were provided with a LENOVO Thinkpad ×200 Tablet (Beijing, China) running Microsoft Windows 7 (Redmond, USA) so that they could practice at home. Participants in the intervention group practiced at home 5 days a week during the 6-weeks long experiment. They were asked to find a quiet environment to practice free of distractions, and, as much as possible, a regular time to practice. The subjects’ performance data was uploaded to a database at the end of each session, so that research assistants could monitor their progression and contact them if they detected any problems with the execution of the task or any lack of improvement, which was rarely necessary.

The sessions at home had two different phases: an unpaced phase (Figure 1 C) followed by a paced phase (Figure 1 D). In the unpaced phase, subjects were instructed to perform the sequence correctly but doing it as fast as possible and attempting to improve their speed continuously. To keep subjects focused, the time to execute the sequence was limited to 5 s. Responses that were incorrect, too slow, or missed (no response) were signaled with a buzzing sound and a red cross mark displayed on screen at the end of the trial, together with a message indicating the reason for the error. If the response was correct, 3 bars were displayed: a blue bar indicated the score (speed, inverse of Movement Time [MT]) of the completed trial, a green bar displayed the subject’s score across past trials and sessions, and a yellow bar displayed a fictitious group best score. Subjects were told that this yellow bar reflected the best score for a group of subjects examined previously and that they should strive to surpass it. To keep the task challenging and at the same time confer a feeling of continuous improvement, the yellow bar was manipulated so that their score got closer and closer with time to the fictitious group score, but they were never able to reach it. This manipulation kept participants motivated throughout experiment and was disclosed at the end of the study. Subjects had to complete 20 trials correctly before moving on to the next sequence, and they did not encounter the same sequence again in the same training session.

In the paced phase participants were supposed to execute the same sequences but following a predefined tempo (0.7 s / beat). After a fixation cross had been shown for 1 s, subjects were prompted to wait during 3 preparation beats that signaled the pace at which to press the keys. This preparation phase was followed by an execution phase of 5 beats in which subjects were supposed to press the sequence of key combinations at the presented tempo. The beats were indicated by a visual counter and an auditory beat. Subjects had to complete 5 trials correctly before moving on to the next sequence. At the end of the trial subjects received feedback regarding whether the response was correct (green tick mark) or incorrect, too fast or too slow (buzzing sound plus red cross and message). Responses were considered too slow (fast) if MT was 10% longer (shorter) than the MT for the prescribed beat. The paced phase was implemented so that subjects could practice the task that was administered during the fMRI measurements, which was paced to rule out that potential differences in neural patterns could be driven by timing differences in the motor output rather than differences in representation. The paced phase in the scanner differed from the one at home in a number of aspects: (1) subjects relied only on the screen counter to follow the tempo and could not hear the auditory beats (because the coil was very narrow and did not allow to use a headset that would have protected their hearing); (2) sequences were interspersed pseudo-randomly (with 2 consecutive trials of the same sequence never taking place), counterbalanced across subjects; (3) no feedback was provided at the end of the trial; (4) instead there was a pseudorandom exponentially distributed inter-trial-interval (ITI) with mean 7.4 s and truncated between 6.0 s and 9.2 s, counterbalanced across subjects; (5) subjects had to perform 2 (of the 3) trained sequences and the same 2 untrained sequences of the unpaced phase; (6) the number of trials was fixed, with 8 trials per sequence and run, yielding a total of 160 trials in a single fMRI session (5 runs). To improve participants’ comfort during the task and avoid that superfluous movements produced undesired BOLD signal fluctuations, after every 8 trials subjects were given 6 seconds to stretch their hand, indicated by the text ‘STRETCH’ displayed on the screen. Supplementary Figure 1 illustrates the timeline of an fMRI task trial. To perform the task in the scanner subjects used an MR-compatible version of the same button box that the experimental group was provided to train at home.

In both the unpaced and paced phase, intervention subjects trained 3 different sequences at home (hereafter *trained sequences* as opposed to *untrained sequences*). Practice sessions at home lasted a median time of 16.3 (SD = 3.7) minutes.

### Motor sequences

To create the sequences, we considered all key combinations (chords) of 4 fingers except 1-1-0-1 and 1-0-1-1 (pinky-ring-middle-index; 1 indicates the finger is used and 0 otherwise). A preceding pilot study showed that these 2 chords were considerably more difficult for subjects and some of them did not manage to press the keys simultaneously after multiple trials. Therefore, they were discarded, leaving 13 different chords to form sequences by concatenation.

We generated sequences of 5 chords with a Hamming distance of 3 between each transition (i.e., 3 fingers had to change between 2 consecutive combinations) which did not contain any repeated chords. These were divided into 2 sets of 3 trained and 6 untrained sequences, which we call configuration A and B, respectively. The sequences forming these two configurations can be seen in Supplementary Table 2. These sets of sequences fulfilled several conditions. First, there were no common transitions between trained and untrained sequences, as we assumed that the core learning component in the task were transitions rather than chords. Second, we maximized the number of chords not shared between trained and untrained sequences, given the previous condition. Third, we tried to match the frequency distribution of the different fingers as much as possible, given the previous two conditions. More details about how the sequences and configurations were obtained can be found in the Supplementary Information.

To guarantee that any possible differences between trained and untrained sequences were independent of the specific sequences trained, 17 out of the 35 subjects in each group received configuration A and the remaining ones received configuration B, meaning that approximately one half of each group trained on different sequences. For tests in the scanner (2 trained / 2 untrained sequences), the A and B configurations were further split into 2 subgroups each (A1/A2/B1/B2), depending on which 2 of the 3 trained sequences the subjects were tested on. Only two sequences were tested inside the scanner at a given session to respect scanning time limits while still achieving enough trials for the individual sequences to ensure adequate power for the statistical analyses.

Our use of the terms *trained* and *untrained sequences* warrants some clarification. Subjects within the same configuration subgroup (A1/A2/B1/B2) were tested every week on the same 3 *trained* sequences, whether they were in the control or the intervention group. The 2 *untrained* sequences tested in each session varied from week to week but were the same for control and intervention subjects if they belonged to the same configuration subgroup. Because there were only 6 *untrained* sequences possible with the constraints explained above, the 2 untrained sequences were cycled every 3 testing sessions, that is, they were the same for testing sessions 0, 3, and 6, the same for 1 and 4, and the same for 2 and 5. Therefore, after testing session 3, untrained sequences were not completely novel, but they had not been seen by subjects for 3 weeks.

To ensure that possible effects of training were not confounded by the order of presentation of sequences in the practice sessions with the laptop, this order was counterbalanced across subjects, and across sessions to minimize potential expectancy effects.

### Testing sessions

Both experimental groups were invited to 7 sessions at the MR center (Figure 1 A). These sessions corresponded to sessions 0 (baseline), 6, 12, 18, 24, 30, 36 for intervention subjects (6 sessions were done within a week, with 5 of practice at home and one of testing at the MR facility). The testing sessions also had 2 phases, with an unpaced phase outside the scanner and a paced phase inside the scanner. The paced task was used in the scanner to avoid that timing differences in the motor output affected activity. The purpose of the unpaced phase was to compare the two groups’ execution speed in laboratory conditions every week on trained and untrained sequences. The unpaced phase in the testing session differed only with that in the home training sessions in that there were 2 additional *untrained* sequences that were tested besides the 3 trained ones. The behavioral tasks always followed structural scanning. Supplementary Table 3 summarizes the characteristics of each phase and session.

### Data collection waves

Because many scanning sessions were required and scanner availability was limited, the data collection was divided into 5 waves of 7 intervention and 7 control subjects each. Due to scanner malfunction, the 4th wave lasted one week less than planned (5 weeks, with 6 scanning sessions). On average, subjects in the intervention group that were included in the 4th wave completed 24.1 (out of 25, SD = 0.9) training sessions at home, whereas those in the remaining waves completed 29.6 (out of 30, SD = 0.8) training sessions.

### Statistical analyses of behavioral data

Subjects in general did not make many errors in a session, but, in a few cases, we observed a disproportionate number of errors (number of errors/session: median = 6, minimum = 0, maximum = 291), in particular for home sessions. We reasoned that the subjects were not paying enough attention or perhaps were experiencing some problem in those sessions. Consequently, we excluded all trials of a particular sequence if the number of incorrect trials for that sequence was more than 1.5 interquartile ranges (IQR) above the 3rd IQR of the number of incorrect trials for any subject for that sequence. This was done separately for each session to allow for a higher number of incorrect trials in initial sessions or more difficult sessions, and led to exclusion of 6.7 % of the data in total.

Only correct trials were included in analyses of MT. Within each session, subjects’ performance was initially not very consistent, and it took a few trials until MT stabilized. We therefore discarded the first 10 trials. With the remainder we computed the median and standard deviation (SD) of MT for each sequence. We also determined the 4 inter-press intervals (IPIs) between the chords that constituted a particular sequence and computed the correlation between IPIs from consecutive trials of the same sequence, averaging the correlation across trials to obtain a measure of execution consistency within a session (Beukema et al. 2019). The same procedure was followed for lags between trials ranging from 1 (i.e., consecutive trials) to 9. Finally, we computed the number of wrong trials that were committed by a subject for each type of sequence and session until reaching the 20 correct trials required.

The 4 dependent variables were analyzed individually and separately for each test session (0-6) with a linear mixed model (LMM) to test for the main effects of group (intervention vs. control group) and practice (trained vs. untrained sequences), and for the interaction between group and practice. A random intercept was included in each model and we covaried for training configuration (i.e., the counterbalanced sets A/B). FDR control (q < 0.05) was applied for the number of sessions. Restricted maximum likelihood (REML) was used to estimate the parameters, with Satterthwaite’s approximation to calculate p-values. These analyses were performed with R Software and the lme4 and lmerTest packages.

### Neuroimaging sessions

The MRI sessions took place at the 7T facility at Skåne University Hospital in Lund (Sweden). Scanning was performed with a 7T Philips Achieva scanner (Best, the Netherlands) with a dual channel transmit, 32-channel receive head coil (Nova Medical, Wilmington, MA, USA). Due to malfunction of the RF system, an 8-channel transmit coil was used in 24 of the sessions.

In each session, a research assistant received the participant at the MR facility, checked MRI eligibility, and ensured that no ferromagnetic objects were brought into the scanner room. Subjects were scanned with earplugs and given a button to press in case they wanted to interrupt the scanning.

To minimize confounds with functionally-related acute differences in cerebral blood flow (Franklin et al. 2013; Månsson et al. 2018), structural scans were always performed before functional scans. Subjects were instructed to avoid practicing the task during the day of the MR scan. In the baseline session, once the structural scans had been acquired, participants were taken outside of the scanner and asked to perform the unpaced part of the task. After completing this part, subjects underwent a functional scan while performing the paced part of the task (see details above). In the remaining sessions, the structural and functional scans were performed consecutively and the unpaced part of the task was done after all scanning had finished. The order was different in the baseline session to give the participants the chance to try out the task before the first functional scan. Otherwise, due to the difficulty of performing the sequences, the error rate during the baseline functional acquisition would have been too high for most participants. Subjects were asked to abstain from alcohol intake on the day before scanning, and prior to the examination on the day of scanning. They were also asked to refrain from caffeine consumption on the day of scanning, as intake of the latter can potentially affect morphometric measures (Ge et al. 2017).

### MRI protocol

In each session, we acquired a T1-weighted (T1w) MP2RAGE scan (Marques et al. 2010), with the following parameters: MP2RAGE_TR_ = 5000 ms, TI_1_/TI_2_ = 900/2750 ms, flip angles = 5° and 3°, TR/TE = 6.8/2.4 ms, 257 sagittal images, matrix size = 320×320, voxel size = 0.7 mm isotropic, SENSE factor = 2, partial Fourier = 75%; the total scan duration was approximately 8 minutes. We also acquired a T2-weighted (T2w) TSE scan, with parameters TR =2500 ms, TE = 314 ms, excitation flip angle = 90°, refocusing flip angle reduced to 35°, 283 sagittal images, matrix size = 320×320, voxel size = 0.7 mm isotropic, SENSE factor = 2×2; the duration of this scan was around 6 min. For functional MRI, EPI scans (TR = 1200 ms, TE = 25 ms, flip angle = 65°, matrix size = 112×116, voxel size = 2×2 mm, 44 axial slices of 2 mm thickness and 0.7 mm spacing, SENSE factor = 3, 5 runs of 400 volumes each) and an auxiliary B0 map were acquired. The total duration of the functional scans was approximately 40 min, excluding short breaks between runs.

### Preprocessing of structural data

Bias-free structural images (also known as *flat* images) were obtained by combining the complex images generated by the MP2RAGE sequence and used to derive CT and GMV. The structural images and surface reconstructions were inspected visually, discarded when not deemed acceptable, and the processing pipelines rerun without them (i.e., the same scans were used for CT and GMV analyses).

#### Cortical thickness (CT)

To produce surface-based maps of CT we processed the structural images with the longitudinal pipeline for FreeSurfer 6.0.1 (https://surfer.nmr.mgh.harvard.edu/), with a modified protocol for skull-stripping the MP2RAGE-derived structural images (Fujimoto et al. 2014). Besides the T1w image, we included the T2w image (-T2pial option) when processing the original images and the average for each subject (within-subject template, also called base image), as the additional contrast can facilitate locating the boundaries of the pial surfaces and lead to better surface reconstructions. For the last step (called long image), to avoid that small differences in geometry or registration between T1w and T2w images affected the results, we only employed the T1w images (see Reuter et al. 2012 for details on the longitudinal processing). The CT maps were registered onto the cortical surface of the average subject’s template (Freesurfer’s *fsaverage*) and smoothed with a kernel with 10 mm of full-width at half-maximum (FWHM) using Connectome Workbench (https://www.humanconnectome.org/software/connectome-workbench). For comparison, the hand-knob area has around 14 mm of diameter (Yousry et al. 1997).

#### Gray matter volume (GMV)

To derive GMV maps, the T1w images were preprocessed with CAT12.7 using the longitudinal preprocessing pipeline (http://www.neuro.uni-jena.de/cat12/), consisting of within-subject longitudinal registration, segmentation into gray matter, white matter and cerebrospinal fluid probability maps, normalization, and smoothing (8 mm FWHM).

### Statistical analyses of structural data

#### Structural Region-of-interest (ROI)

In order to reduce the number of tests in the univariate analyses, we created and preregistered a mask encompassing cortical areas involved in motor sequence learning (Wiestler and Diedrichsen 2013; Yokoi 2019; Berlot et al. 2020), where changes related to training the motor task were expected to take place. For this purpose we aggregated a number of frontal and parietal parcels from the Human Connectome Project’s Multi-modal Cortical Parcellation (Glasser et al. 2016). Supplementary Figure 8 shows the mask, and a list with the parcels that were combined can be found in Supplementary Table 4. This mask (henceforth ROI_surf_) was created on the fsaverage surface, and transferred to MNI space to create a mask for volumetric analyses (henceforth ROI_vol_).

#### Reliability analyses

At each vertex/voxel we computed the intraclass correlation coefficient (ICC) separately for CT and GMV, using a two-way random-effects model to estimate agreement across timepoints with the *irr* R package. A high ICC at a given voxel indicates that the value of the measure of interest at that voxel has low within-subject variance across timepoints, compared to the between-subject variance.

#### Univariate structural analyses

We used linear mixed models (LMMs) with the fixed effects of experimental group (intervention vs. control; coded as a factor), test session (linear and/or non-linear; see below), and the group x session interactions to test for effects of training on brain structure (predicting the interaction effects). Separate models were estimated with CT and GMV as dependent variables. Only random intercepts were included in the models, since including random effects for the linear and/or quadratic slopes (i.e., the effects of test session) often led to singular models (variance close to 0) and their inclusion made no meaningful difference for the fixed effects estimates we were interested in. This holds also for the other analyses below involving LMMs.

At each vertex/voxel we computed the value of the Bayesian Information Criterion (BIC) for 5 different LMMs that differed in how session was specified: (1) a model with only a linear term for session interacting with group; (2) a model with an asymptotic term for session interacting with group (increasing inverse-quadratically until the middle of the training period and constant afterwards); (3) a model with an inverse-quadratic term of session interacting with group; (4) a model with linear and quadratic terms of session interacting with group; and (5) a model with linear, quadratic and cubic terms of session interacting with group. All these models had, additionally, linear, quadratic and cubic terms for session (main effects). Models 1-3 were preregistered (Supplementary Figure 9), although the cubic-polynomial main effects were not considered initially and were added a posteriori upon observation of the data, to account for non-linear drifts in structural measures over time in the whole sample, which could compromise the ability to detect interaction effects. Models 4 and 5 were also added post hoc for completeness in case that models 1-3 were too rigid to fit the data adequately. Note that models 4 and 5 incorporate, respectively, 1 and 2 more parameters than models 1-3, but this increased flexibility should be penalized by the BIC.

With the winning models, we tested for interactions between experimental group and session terms. P-values for interaction terms and for tests considering collectively for the different session terms (e.g., for model 4, linear and quadratic terms) were obtained with the Type III Wald chi-square test (as implemented in the Anova function of the *car* R package).

The univariate tests were restricted to the vertices in ROI_surf_ for the analysis of CT, and to the voxels in ROI_vol_ for the analysis of GMV. Since in a few sessions the regular transmit coil had to be replaced by an auxiliary one due to malfunctioning, we introduced this covariate of no interest in the model. We also controlled for training configuration and framewise displacement (FD, Power et al. 2012) estimated during the functional scan to approximate in-scanner motion during the structural scan, since its potential impact on morphometric measures has been documented (Reuter et al. 2015).

### Preprocessing of functional data

Functional MRI data were preprocessed with fMRIPrep version 20.1.1 (Esteban et al. 2019), a Nipype-based tool (Gorgolewski et al. 2011). The corresponding preprocessing steps are described here using the citation boilerplate provided by the software. Each T1w volume was corrected for intensity non-uniformity using N4BiasFieldCorrection v2.1.0 (Tustison et al. 2010) and skull-stripped using antsBrainExtraction.sh v2.1.0 (using the OASIS template). The brain mask estimated previously was refined with a custom variation of the method to reconcile ANTs-derived and FreeSurfer-derived segmentations of the cortical gray-matter of Mindboggle (Klein et al. 2017). Spatial normalization to the ICBM 152 Non-linear Asymmetrical template version 2009c (Fonov et al. 2012) was performed through non-linear registration with the antsRegistration tool of ANTs v2.1.0 (Avants et al. 2008), using brain-extracted versions of both T1w volume and template. Brain tissue segmentation of cerebrospinal fluid (CSF), white-matter (WM) and gray-matter (GM) was performed on the brain-extracted T1w using FAST (Zhang et al. 2001). The subject-average cortical surfaces reconstructed with the longitudinal pipeline for structural analyses were also employed for surface-based functional analyses.

Functional data were motion corrected using mcflirt (FSL v5.0.9, Jenkinson et al. 2002). Distortion correction were performed using fieldmaps processed with FUGUE (FSL v5.0.9, Jenkinson 2003). This was followed by co-registration to the corresponding T1w with 6 degrees of freedom with FLIRT (FSL v5.0.9, Jenkinson 2003). We did not use boundary-based registration because in our 7T functional data, it led to failed registrations in a considerable number of cases, whereas the volume-based method yielded accurate registrations. Motion correcting transformations, field distortion correcting warp, BOLD-to-T1w transformation and T1w-to-template (MNI) warp were concatenated and applied in a single step using antsApplyTransforms (ANTs v2.1.0) using Lanczos interpolation. Functional runs with excessive head motion (FD more than 1.5 times the interquartile range above the upper quartile) were removed (3.6% of runs).

### Statistical analyses of functional data

#### Functional ROIs

We defined 4 ROIs in each hemisphere (Figure 4 A), encompassing cortical areas involved in motor sequence learning (Wiestler and Diedrichsen 2013; Yokoi 2019; Berlot et al. 2020), by aggregating parcels from the Human Connectome Project’s Multi-modal Cortical Parcellation (Glasser et al. 2016):

Primary sensorimotor (PS): R_1_ROI, R_3a_ROI, R_3b_ROI, R_4_ROI (right hemisphere); L_1_ROI, L_3a_ROI, L_3b_ROI, L_4_ROI (left hemisphere), excluding vertices further than 25 mm of distance from the respective hand knob (Yousry et al. 1997; Berlot et al. 2020).

Premotor (PM): R_FEF_ROI, R_6a_ROI (right hemisphere); L_FEF_ROI, L_6a_ROI (left hemisphere).

Supplementary motor area (SMA): R_6mp_ROI, R_6ma_ROI, R_SCEF_ROI (right hemisphere); L_6mp_ROI, L_6ma_ROI, L_SCEF_ROI (left hemisphere).

Superior parietal lobule (SPL): R_AIP_ROI, R_IP2_ROI (right hemisphere); L_AIP_ROI, L_IP2_ROI (left hemisphere).

#### Activation analyses

The preprocessed fMRI data were resampled on the fsaverage6 surface (with 41k vertices for each hemisphere), and a smoothing kernel with 10 mm FWHM was applied with Connectome Workbench. The smoothed data was analyzed with the FMRIB’s Software Library (FSL, https://fsl.fmrib.ox.ac.uk/fsl/fslwiki/). Low-frequency drifts in the time domain were removed by applying a high-pass filter cutoff of 90 s. In the single-subject-level statistical analyses, the general linear model (GLM) matrix included regressors formed as boxcar functions convolved with double-gamma hemodynamic response functions accounting respectively for trained correct trials, trained incorrect trials, untrained correct trials, untrained incorrect trials and fixation and hand-stretching periods. Six realignment parameters to correct for head motion plus the derivatives of these and all the previous regressors were also included in the model.

#### Univariate activation analyses

On the group-level, we used LMMs to test for effects of motor training on brain functional activation. The parameter estimates from the first level analyses corresponding to activation relative to baseline were used as the dependent variable. The model included, for each subject, a random intercept and linear and/or non-linear fixed effects of session number. At each vertex/voxel in the 4 functional ROIs we computed the BIC to compare the same 5 LMMs explained for the structural analyses above (linear, asymptotic, quadratic, quadratic plus linear and cubic), with the only difference that the interactions of session terms in this case were not only with group but also with practice (trained/untrained), both coded as factors. We tested for group x session terms, practice x session terms and group x practice x session interactions with the winning model for each vertex.

The univariate tests were restricted to the vertices in a mask formed by the 8 cortical ROIs (4 per hemisphere) mentioned above. False discovery rate (FDR) control for multiple testing across voxels was employed, with q < 0.025 (to account for the two hemispheres). Like for structural analyses, to rule out the influence of the type of transmit coil used or of motor training configuration, we introduced these covariates of no interest in the models. We also controlled for average framewise displacement within each run to account for in-scanner motion.

#### Multivariate pattern analyses of variability

For each voxel and functional run, we first estimated trial-by-trial activation patterns (Mumford et al. 2012) by fitting to each voxel’s data a GLM model with one regressor per trial (impulse response convolved with a double gamma hemodynamic response function), plus 2 additional regressors accounting respectively for fixation and hand-stretching periods. Six realignment parameters to correct for head motion plus the derivatives of these and all the previous regressors were also included in the model. No smoothing was applied to the data to estimate these parametric maps. To take into account that voxel signal is corrupted by noise, we applied multivariate spatial prewhitening (Walther et al. 2016) of the regression coefficients with a regularized estimator of the variance-covariance matrix of the residuals (Ledoit and Wolf 2004).

Next, we were interested in quantifying the variability of these trial-by-trial activation patterns separately for trained and untrained sequences, so as to be able to compare them. Given the set of voxels *R* in one of the cortical ROIs, and *V*_*t*_ = {*β*_*i,t*_, *i ∈ R*}, the activation pattern in the ROI (vector of parameter estimates) for trial *t*, we calculated the trial-by-trial matrix *G* formed by the scaled inner products of activation patterns, 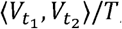, for all trial pairs (*t*_1_, *t*_2_)in a session, which is related to their similarity. For this estimation we included in our analyses correct trials only, and only from runs with at least 3 correct trials for each of the 4 sequences tested. The matrix *G* determines the representational geometry of activity profiles, and Euclidean or cosine distances can be easily derived from it (Diedrichsen and Kriegeskorte 2017; Diedrichsen et al. 2017). We then expressed *G* as a linear combination of 8 components specifying the contribution to its structure from different features related to the similarity between sequence pairs:

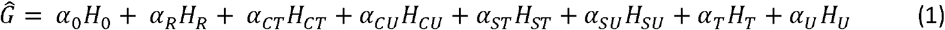

*H*_*o*_ corresponds to a global intercept (matrix with all entries equal to 1). 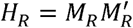 models the increased covariance for pairs of patterns of the same run, where *M*_*R*_ is an indicator matrix with a dummy variable for each run. *H*_*CT*_ reflects the similarity for pairs of patterns of trained sequences, shared across runs, with 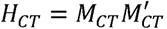 and *M*_*CT*_ an indicator vector with ones for trained sequences and zeros otherwise. *H*_*CU*_ is equivalent to *H*_*CT*_ for untrained sequences. 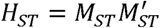 reflects the similarity for trained sequences of the same type (in a session, 4 different types of sequences were presented, 2 of which were trained sequences and 2 untrained sequences, and each of these sequence types was executed 40 times), with *M*_*ST*_ an indicator matrix with a dummy variable for each type of trained sequence, with ones when the trial corresponded to that type and zero otherwise. *H*_*SU*_ is equivalent to *H*_*ST*_, for untrained sequences. Finally, *H*_*T*_ is a diagonal matrix where the diagonal is an indicator vector with ones for trials for trained sequences, and similarly for *H*_*U*_. Thus, the coefficients for *H*_*T*_ and *H*_*U*_ should reflect the variability for trained and untrained sequences, respectively.

Since the coefficients *α* need to be positive, we used non-negative least squares to estimate them. The coefficients of interest are *α*_*T*_ and *α*_*U*_ as they reflect the magnitude of the variability of trained and untrained sequences respectively, together with fMRI noise. Under the assumption that the level of noise for both types of sequences is comparable, we can use these coefficients and *α*_*U*_ and *α*_*U*_ to compare the variability of trained vs. untrained sequence patterns, taking logarithms to render the estimates normally distributed for successive analyses. Finally, we computed the variability index *s = log (α)* for each session, subject, sequence type group/untrained) and each of the 8 cortical ROIs specified above. These calculations were implemented in Python 3.6.

#### Dissimilarity of neural patterns

To elucidate further the changes in neural patterns over the course of the experiment, for each of the 8 cortical ROIs we computed one further metric, the squared cross-validated Mahalanobis distance (also known as cross-nobis dissimilarity (Walther et al. 2016; Nili et al. 2020)) between patterns of different sequence type:

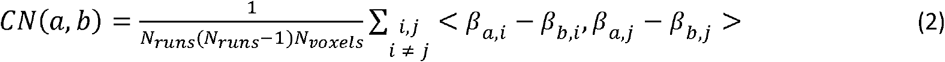

where <·,·> denotes scalar product; *a, b* index sequence types; *i, j* index different runs and *β*_*a,i*_, is the average of trial-wise patterns of type *a* within run *i*. We used a cross-validated metric because its non-cross-validated counterpart tends to be biased by the presence of noise (Walther et al. 2016; Diedrichsen et al. 2021). The cross-validation folds corresponded to the different session runs, as the noise across different runs can be considered independent. As seen in equation (2), to obtain the metric we divided the summed scalar products by the number of voxels *N*_*voxels*_, rendering them comparable across subjects. We arranged the ensuing metric estimates by whether the pair of sequences was formed by two trained, two untrained, or by one trained and one untrained sequence. Then we averaged the estimates within each of these categories.

To further illustrate the changes in neural patterns, we also trained a linear support vector machine (SVM) to discriminate between patterns from different pairs of sequence types. Like for the previous metrics we arranged the pairs when averaging to calculate session/subject estimates by whether they were from trained, untrained or both trained and untrained sequences. Classification accuracies were estimated by cross-validating across runs, and the SVM regularization parameter C was optimized using nested cross-validation. These calculations were implemented in Python 3.6.

#### Statistical analyses of dissimilarity and variability of neural patterns

For cross-nobis dissimilarities, at each session, we tested for a group x practice interaction using an LMM, where in this case the practice factor refers to whether the dissimilarity is between one trained and one untrained sequence or between different untrained sequences. For the variability index (*s* = *log* (*α*) and the variance of neural patterns, we tested for a group x practice (trained/untrained) interaction at each session, separately for each session. In these analyses, we applied FDR control for the 8 ROIs and number of sessions, with q < 0.05. To rule out the influence of the type of transmit coil used, motor training configuration or in-scanner motion, we introduced these covariates of no interest in the models.

### Datasets used for different analyses

As noted above, data collection was divided in 5 waves of 14 subjects each for logistic reasons. The first wave was used for the purpose of piloting (although the protocol was not changed in subsequent waves and the data should be equivalent). Thus, the preregistered hypotheses need to be confirmed using data from the last 4 waves only. For analyses where we were able to confirm the preregistered hypothesis, we report results for the 4 waves. Whenever the results were either negative or the analyses were not preregistered, there was no reason to exclude the first wave and therefore we report results corresponding to the full dataset.

### Data/code availability

The European General Data Protection Regulation (GDPR) and supplementary Swedish data protection legislation prohibit us from making the data publicly available, but data can be requested and subsequently transferred for projects with well-defined analyses (described in a project outline) that have been approved by the requesting researcher’s local ethics committee and are consistent with the original ethics approval. This requires a data sharing agreement that effectively transfers the confidentiality obligations of the host institution where the original research was carried out to the institution receiving the data.

The code is available from the following repositories: https://github.com/benjamingarzon/SeqLearn (motor task) https://github.com/benjamingarzon/LH-RSA (analyses)

## Results

### Behavioral measures

In this section we analyze the data of the unpaced task, where subjects were asked to execute the sequences correctly but as fast as possible, always trying to improve their speed. We report the pattern of behavioral changes as a function of experimental group (intervention vs. control) and sequence type (trained vs. untrained).

We first asked whether the intervention subjects improved on the speed of executing the sequences that were trained at home by this group 5 times a week over 6 weeks. This was the case. Median movement time (MT) displayed rapid initial reductions that eventually stabilized (Figure 2 A). The learning curves were reliable in the sense of being more similar within (i.e., across trained sequences) than between subjects (see Supplementary Information).

**Figure 2.**
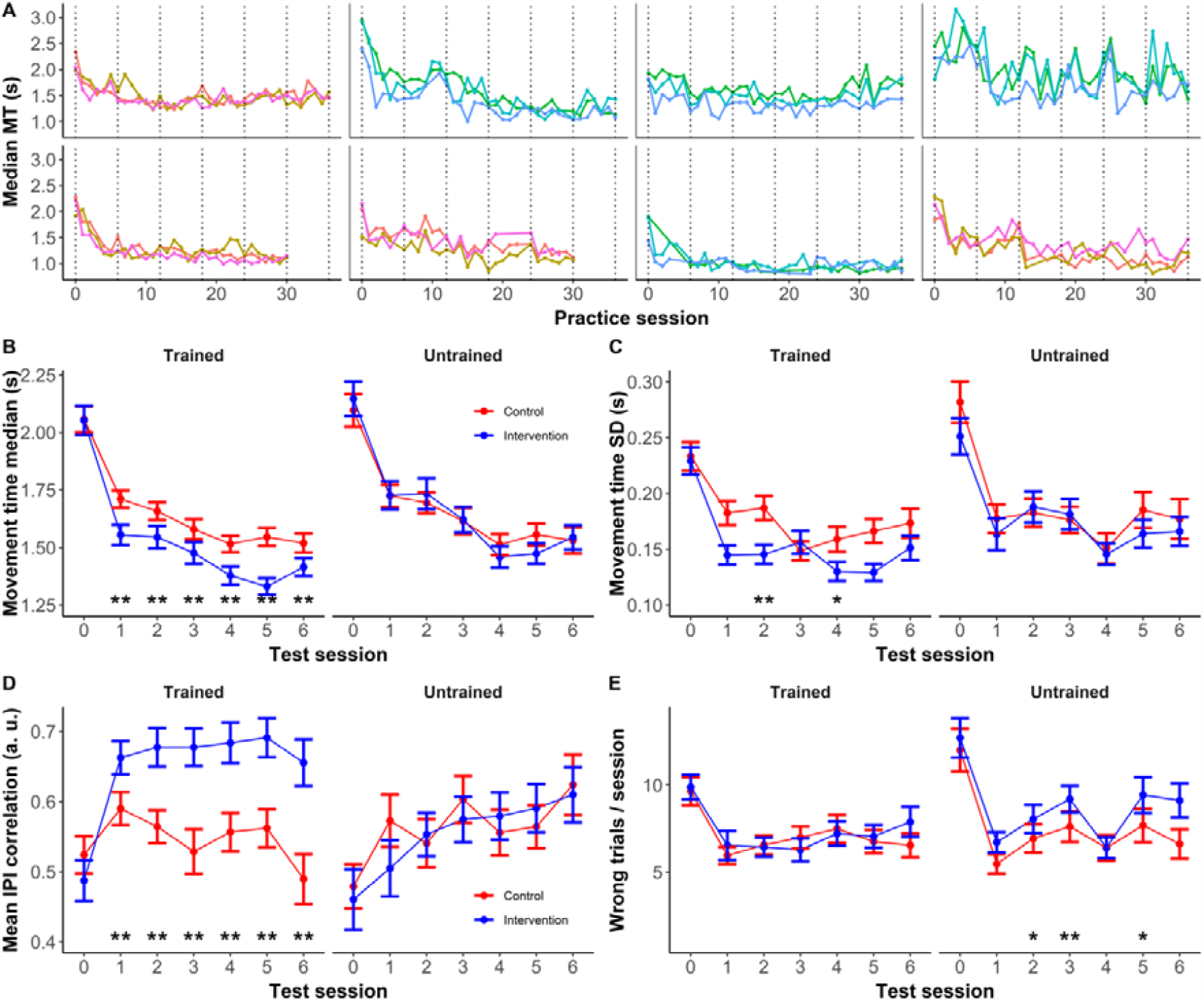
Behavioral measures. A) Median movement time (MT) over practice sessions for 8 representative intervention participants showing reductions in MT with training and interindividual variability in the learning curves. Different trace colors denote different sequences trained. The dotted vertical lines correspond to the 7 on-site test sessions, referenced in the remaining panels. B) Mean of the median (within-session) MT for the two groups and sequence types (trained/untrained). C) Standard deviation of MT, as a measure of performance variability. D) Correlation between the inter-press intervals (IPIs) of consecutive trials, a measure of performance consistency. E) Number of incorrect trials in a session. In all panels, the measures for the control group are depicted in red, and for the intervention group in blue. The measures are averaged across subjects and specific sequences. Error bars denote standard error of the mean. Where indicated, the interaction of group x practice (trained/untrained) was significant: ^‘**’^ significant FDR-corrected, q < 0.05; ^‘*’^ significant uncorrected, p < 0.05.

Because the intervention subjects and the control subjects (who did not train on the sequences) were both tested in the lab once a week on the 3 sequences that the intervention group trained at home (trained sequences) and 2 additional ones (untrained sequences), we could also address whether there were training-related sequence-specific and sequence-general improvements, and whether there were global effects of repeated testing behind the improvements over time. The overall time-course of MT followed a differentially faster decrease for the intervention group and the trained sequences relative to the control group and the untrained sequences (Figure 2 B). The group (intervention vs. control) x practice (trained vs. untrained sequences) interaction reached significance for every testing session after baseline, reflecting that intervention subjects were faster than controls especially on trained relative to untrained sequences after a week of training (p < 0.05, FDR-corrected for the number of sessions; see Supplementary Table 5 for the statistics in each session). Thus, parts of the performance improvements displayed by intervention subjects were specific to the trained sequences. In addition to these sequence-specific improvements, both groups showed substantial improvements in median MT over the training period, with most of the decreases taking place initially; the intervention subjects did not show significantly larger improvements on untrained sequences than the control participants (Figure 2 B). Overall, this pattern of results suggests that there was no detectable transfer of learned improvements to novel sequences. The improvements for the untrained sequences also occurred for the control group, and therefore were due to the learning occurring in the test phases, and may contain both sequence-specific and sequence nonspecific improvements. Such aspects may relate to goal selection, online planning, and the use of explicit knowledge to find a general strategy to perform the task (Krakauer et al. 2019; Spampinato and Celnik 2020; Ariani et al. 2021).

Because past studies have found reductions in behavioral variability over learning, we also investigated whether training reduced the variability of motor output. Both groups showed decreases in variability (SD) of MT for both sequence types, with a larger reduction over time for trained sequences in intervention subjects relative to controls and untrained sequences (Figure 2 C). The interaction group x practice was statistically significant (p < 0.05, FDR-corrected for the number of sessions) in test session 2 (and significant uncorrected in test session 4; see Supplementary Table 5 for statistics). To provide a complementary measure of behavioral variability, we also correlated the IPI pattern (i.e., the 4 intervals between the 5 discrete presses in a sequence) across trials. Participants in the intervention group demonstrated more highly correlated IPIs between consecutive trials of the same sequence type compared to baseline for trained sequences relative to the controls and to the untrained sequences (Figure 2 D). The group x practice interaction was statistically significant (p < 0.05, FDR-corrected for the number of sessions) in all test sessions after baseline (see Supplementary Table 5 for statistics). When computing the correlation for pairs of trials lagged by several trials the correlation was lower the further apart the trials were, but a similar pattern was observed, namely that practicing the trained sequences led to a markedly more consistent performance on those specific sequences (Supplementary Figure 2). Thus, training reduced execution variability and this effect was partly specific for the trained sequences.

The number of incorrect trials per session and sequence was similar for intervention and control subjects for trained sequences, that is, subjects did not seem to trade-off speed for accuracy when learning the sequences (Figure 2 E). Surprisingly, intervention subjects who practiced at home committed more errors on untrained sequences than the control subjects. The interaction group x practice was statistically significant (p < 0.05, FDR-corrected for the number of sessions) in test session 3 after baseline and uncorrected in sessions 2 and 5 (see Supplementary Table 5 for statistics). These results may indicate that intervention subjects became less careful over time when facing new sequences.

Taken together, these results show both task-general performance improvements and increased performance consistency independent of group or sequence type, but also differential and specific improvements to the sequences that were trained intensively.

Finally, there were considerable differences between trained and untrained sequences for both groups at baseline in MT variability (untrained > trained, t(220.14) = 2.9, p = 0.003, Figure 2 C) and number of correct trials (untrained > trained, z = 2.8, p = 0.006, Figure 2 E), which reflects differences in the difficulty of the two sets of sequences (due to the different cardinality of the sets and the constraints we imposed on the sequences, it was not possible to counterbalance them completely, only to make the proportions of training configurations equivalent for the two groups, see Methods). These baseline behavioral differences are, however, unlikely to account for group differences in differential changes over time. In addition, the statistical analyses included training configuration as a covariate. Therefore, the baseline differences between the trained and untrained sequences should neither affect the main interpretations of the behavioral results nor the imaging results reported below.

### Training-related changes in functional activation

Initial analyses of the fMRI data focused on the activity elicited by task performance. Whole-brain surface-based univariate analyses of the BOLD signal (correct execution > resting baseline), demonstrated task-related functional activation (p < 0.025, FDR-corrected) in bilateral secondary motor areas (supplementary motor area, superior parietal and premotor cortex), primary motor and somatosensory regions, most prominently on the right side, i.e. contralateral to the hand used to perform the movements, as well as in primary visual cortex and regions of the salience network (insular and anterior cingulate cortices; Figure 3 A). Our task thus elicited activity in the expected sensorimotor and dorsal attention networks.

**Figure 3.**
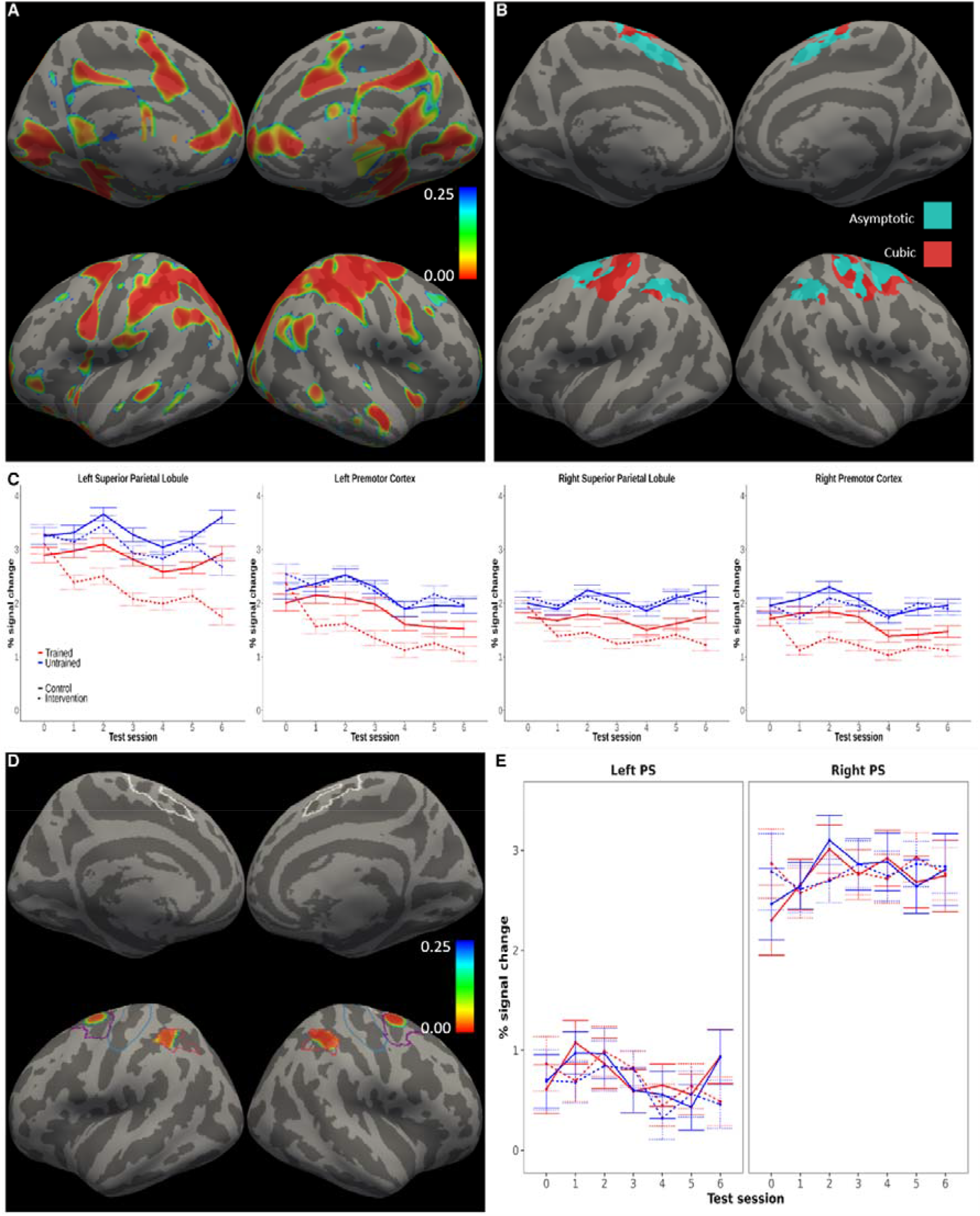
Functional activation. Practice-related changes in functional activation within cortical motor areas. A) The figure shows the significance map (p-values) for the contrast of activation against resting baseline (mean effect across subject groups, timepoints and sequence types). P-values were FDR-corrected considering the whole cortex as the search area. Executing the motor sequences required by the task elicited brain activity in primary and secondary motor regions. B) Result of the model comparison indicating, at each vertex, the model with the lowest BIC (cyan = asymptotic, red = cubic). Non-linear models, and the asymptotic regime in particular, were preferred in the major part of the cortical areas probed. BIC reflects the likelihood of the model penalized by its complexity. C) Functional activation time-courses from clusters where the univariate analyses identified effects of practice. D) Practice x session effect for the asymptotic model, revealing changes over time in activation in bilateral parietal and premotor regions that were differential for trained as compared to untrained sequences. The figure shows p-values FDR-corrected within the preregistered areas for the test of practice-related effects. The pattern found when fitting the cubic model was equivalent. The contour lines mark the preregistered mask encompassing primary sensorimotor and secondary motor cortical areas, and the corresponding analyses were restricted to these areas. E) Average of the functional activation time-courses within the primary sensorimotor (PS) ROIs, in which no significant practice x session effects were detected. In (A, D), the corrected significance threshold was set to *q* = 0.025 to account for both hemispheres.

Next, we tested the preregistered hypothesis of training-related decreases in activation. For these analyses, we performed voxel-wise analyses restricted to a preregistered mask encompassing primary and secondary cortical sensorimotor regions. The analyses showed sequence-specific decreases in activity in secondary sensorimotor areas, but not in primary areas, for the intervention group and the trained sequences relative to the control group and the untrained sequences (Figure 3 C, D). In the analyses revealing these results, we fitted several models that assumed an interaction between experimental group, practice (i.e., trained vs. untrained sequences), and different shapes of changes over the sessions (i.e., the 7 longitudinal measurements; see Methods). To evaluate the fit of these models, we computed the BIC at each voxel in the mask. Within the cortical areas of interest, the BIC was lowest for either the asymptotic or the cubic model, depending on the specific region (Figure 3 B). The practice by session interaction reached statistical significance (p < 0.025, FDR-corrected) with both models in clusters within bilateral superior parietal and premotor cortices (Figure 3 D; Table 1). However, we were unable to detect practice-related effects in primary sensorimotor cortex (no interactions concerning effects of practice were significant in this region even at the uncorrected level). When plotting the effects (Figure 3 C, E), the pattern of results was in line with our preregistered hypothesis only in the secondary sensorimotor areas, which showed larger reductions in activity over time for trained than untrained sequences especially for the intervention group. Note also that control analyses with the linear or quadratic models of the time-trends over sessions did not reveal any additional clusters showing statistically significant practice-related effects. The predicted three-way interaction of group by practice by session approached statistical significance in the same regions reported above for the asymptotic model, but the effect did not survive correction for multiple comparisons. This pattern of results should be interpreted considering the study procedures, which included testing control subjects on a subset of sequences (trained sequences) more often than on the remaining ones, namely every week. This may explain the trends for activation decreases that can be observed also in the control group for the trained set compared to the untrained set, which varied from week to week.

**Table 1.**
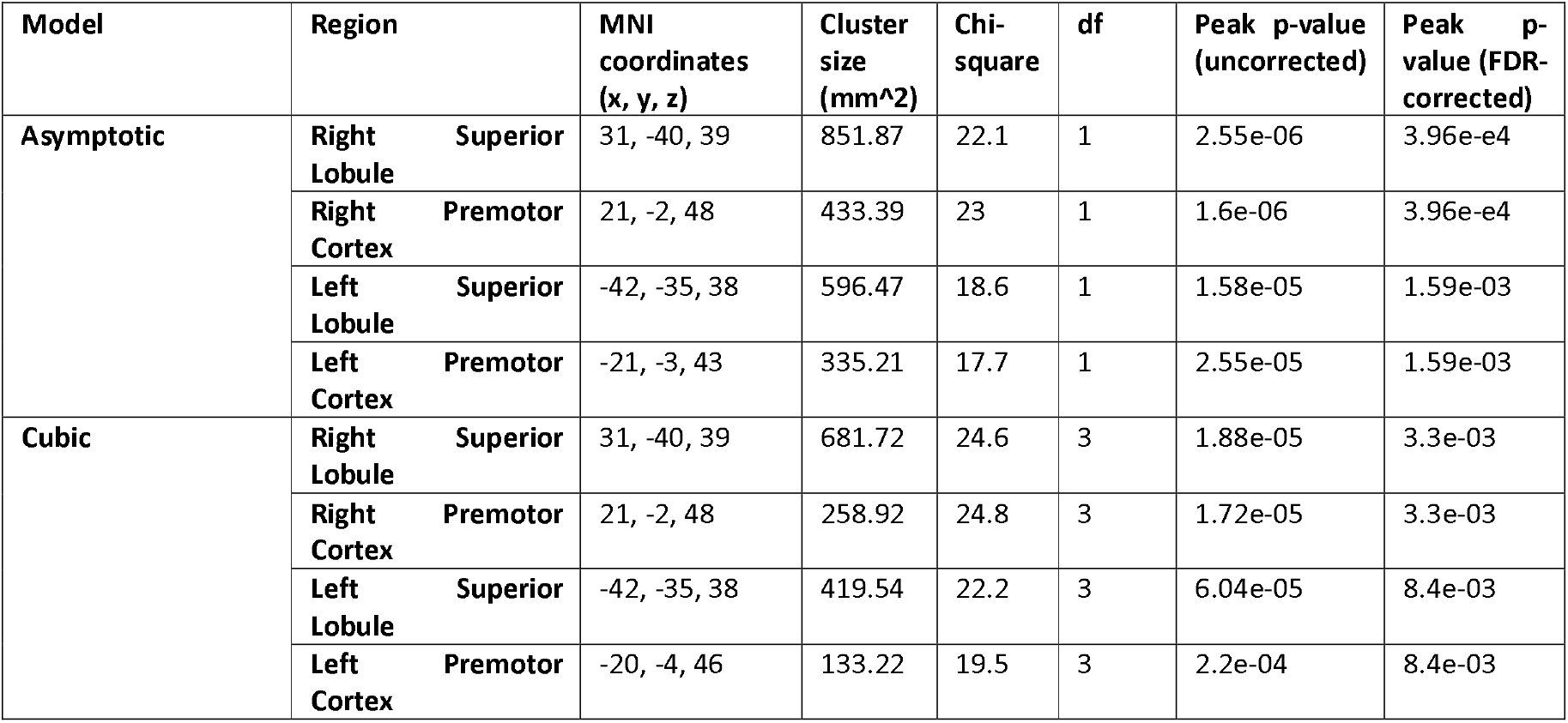
Effects of practice on functional activation. Clusters showing a practice (trained/untrained) x session interaction on functional activation (cf. Figure 3).

Because our design included a control group, we could also test the hypothesis that training effects on activation would generalize to untrained sequences, such that decreases in activity would also be observed in the intervention group relative to the control group for untrained sequences. The results did not support this hypothesis (Figure 3 C). The group by session interactions did not reach statistical significance even at more liberal statistical thresholds. This was confirmed by follow-up tests: when restricting the analysis of the data to untrained sequences, there was no statistically significant differences between groups (p > 0.1 in all clusters). Therefore, practicing the trained sequences resulted in no noticeable activation changes for the untrained ones.

In summary, brain activity for trained sequences decreased relative to untrained sequences in the bilateral parietal and premotor cortices. Training-related changes in the primary sensorimotor areas were not detected.

### Training-related changes in variability of activation patterns over repeated trials of trained sequences

We had preregistered the hypothesis that training specific sequences would result in lower variability of multivariate activation patterns among trials of those sequences, as predicted by the ESR model due to stabilization of neural circuits in its refinement phase. To test this hypothesis, we derived an index of the variability of the multivariate activity patterns over repeated trials of trained and untrained sequences for each subject (see Methods) and examined the group averages over time in preregistered ROIs in SPL, SMA, PM, and PS in each hemisphere (Figure 4 A; these ROIs together formed the mask used for the univariate analyses above). This index of neural variability was not affected by training in a significant manner (Supplementary Figure 5 and Supplementary Table 7; p > 0.1 in all ROIs and sessions, testing separately for a group x practice interaction in each ROI/session and applying FDR control for sessions and ROIs).

**Figure 4.**
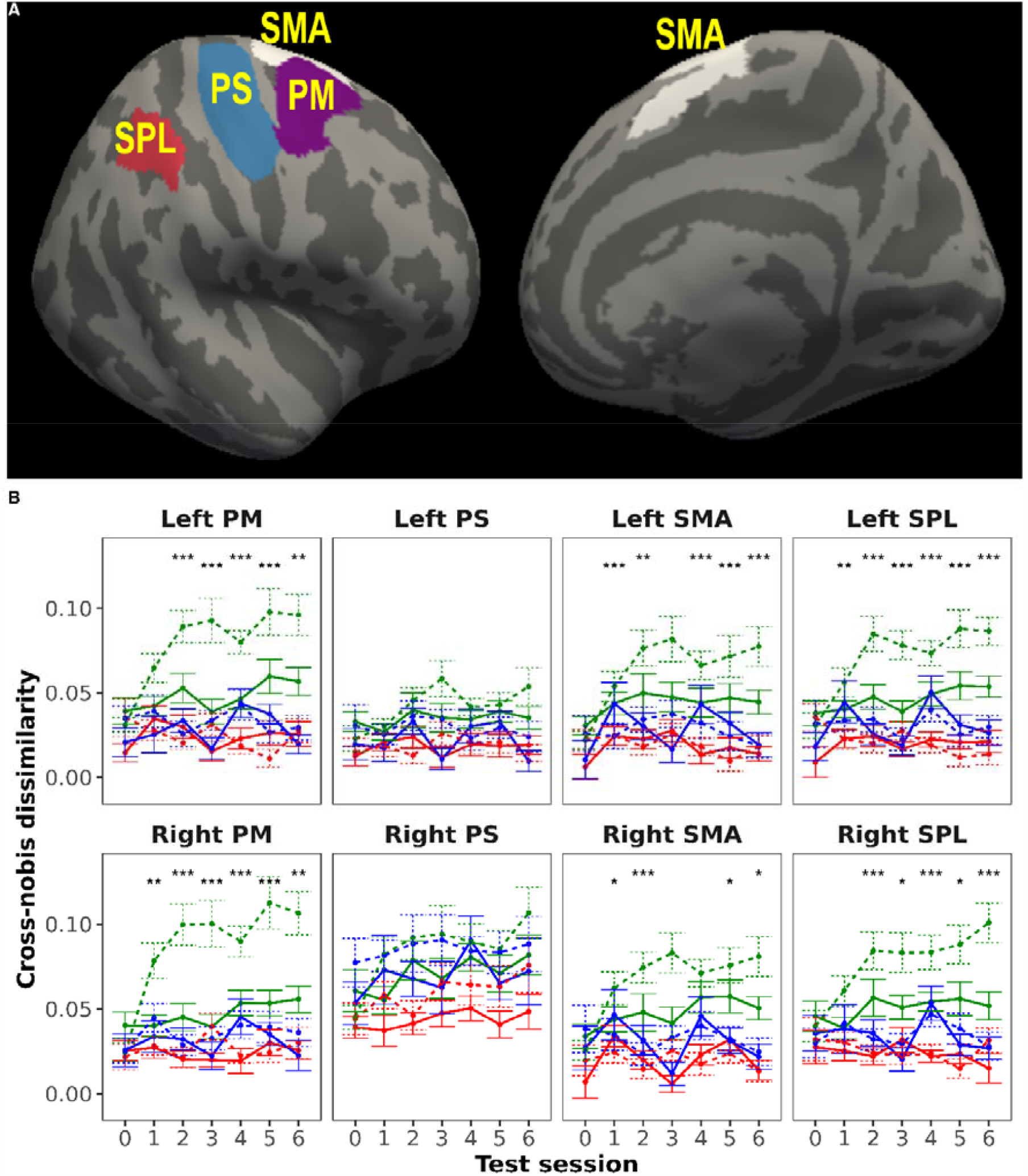
Evolution of cross-nobis dissimilarities between neural patterns for pairs of sequences. A) ROIs that were used for neural pattern dissimilarity analyses (only right hemisphere regions are shown, but the same regions from the left hemisphere were also analyzed). B) Cross-nobis dissimilarities between multivariate patterns of trained and untrained sequences increased over time in both groups, and much more prominently for the intervention group. These changes were present in all regions except PS. Asterisks indicate a significant interaction of group x practice (untrained/trained - untrained). ROI: region-of-interest; PM: premotor; PS: primary sensorimotor; SMA: supplementary motor area; SPL: superior parietal lobule; ^‘***’^ significant FDR-corrected for ROIs and sessions, q < 0.05; ^‘**’^ significant FDR-corrected for sessions, q< 0.05; ^‘*’^ significant uncorrected, p < 0.05.

### Training-related changes in the dissimilarities between activation patterns

To investigate whether multivariate activity patterns, just like the overall activity, also displayed sequence-specific changes with learning, we calculated cross-nobis dissimilarities as a measure of the dissimilarity of activation patterns within and between trained and untrained sequences (see Methods; these analyses were not preregistered). Dissimilarities were computed for the ROIs in SPL, SMA, PM, and PS in each hemisphere (Figure 4 A) as in the previous analyses of pattern variability. The cross-nobis dissimilarities between patterns for trained and untrained sequences increased over time in both groups in all ROIs except bilateral primary sensorimotor areas (Figure 4B, green dashed trace). Nevertheless, the increase was more prominent for the intervention participants and dissimilarities between trained and untrained sequence patterns relative to dissimilarities between untrained sequence patterns and controls (the interaction was significant in several sessions after baseline; Figure 4 B; Supplementary Table 6). In contrast, dissimilarities between sequence patterns for different trained sequences (red traces) or between sequence patterns of different untrained sequences (blue traces) showed generally a stable trend over time in both groups. Overall, these results indicate that practicing the trained sequences resulted in their neural patterns becoming more dissimilar to those from untrained sequences in the secondary motor regions that had shown the largest activation decreases.

The use of the cross-nobis dissimilarities is closely related to the multivariate pattern classification approach that is more common in the fMRI literature. To make a direct contact with this literature and further illustrate our findings, we also report average cross-validated classification accuracies for multivariate models (SVM) trained to separate pairs of different sequences (see Methods). These accuracies were significantly above chance (i.e., accuracy = 0.5) for untrained sequences in all regions in the control group (p < 0.05 in all ROIs for the group mean, considering all sessions), implying that different sequences had to some extent distinguishable neural representations even when they were not practiced (Supplementary Figure 6 A; see permutated data for comparison in Supplementary Figure 6 B). In line with results from computing the cross-nobis dissimilarities, accuracy of the classification between patterns of trained and untrained sequences increased over sessions and more highly for the intervention group (Supplementary Table 8), but remained stable for classification between different trained or untrained sequences.

In summary, the results from these non-preregistered analyses were aligned with the overall activation results by showing that when subjects practiced certain motor sequences the corresponding activation patterns in secondary, but not primary, motor areas became more differentiated from those of untrained sequences. The similarity of the activation patterns among the trained sequences did not show reliable change.

### Structural imaging analyses

To test the preregistered hypothesis of practice-dependent changes in cortical thickness and gray matter volume, we fitted several models that assumed an interaction between experimental group and different shapes of changes over the sessions (i.e., the 7 measurements; see Methods). To evaluate the fit of these models, we computed the BIC at each vertex/voxel of a preregistered ROI. Depending on the region, the BIC was minimized by either the linear, the asymptotic, or the quadratic model, but there was no anatomical congruency (i.e. no clear spatial pattern) in the spatial BIC map. We then tested for interactions between group and session terms within the ROI, but we could not find any clusters in which these tests survived correction for multiple comparisons for any of the measures, even using a liberal threshold. Although there were clusters at the uncorrected level, these were close to the significance threshold and examination of the corresponding time-courses suggested that they were driven by fluctuations in the measures at a few timepoints. That is, we could not identify any pattern consistent with gradual differential increases, decreases, or non-linear progression in the intervention group relative to the control group.

Note that the estimates of cortical thickness and gray matter volume displayed acceptable reliability across timepoints. Intra-class correlation coefficients (ICC; computed separately for each vertex/voxel within the ROI, including only the control subjects and considering these as the class of interest) of the structural measures ranged from moderate to good across vertices (Koo and Li 2016) for CT (median = 0.78, SD = 0.09) and good to excellent across voxels for GMV (median = 0.92, SD = 0.11; Supplementary Figure 3).

In summary, despite the acceptable reliability of the structural measures, we were unable to detect training-related changes in any of the measures. Preregistered supplementary analyses on approximate T1 relaxation times derived from the structural scans, which exhibited lower reliability than morphometric measures, did not reveal any experience-dependent changes either (see Supplementary Information).

## Discussion

In this study, we acquired repeated behavioral performance and neuroimaging measures over the course of 6 weeks to investigate neural changes associated with motor sequence learning. Both the intervention group and control groups showed general performance improvements, but performance improved more, and became more consistent, for sequences that were intensively trained by the intervention group relative to those that were not. In line with our preregistered hypothesis, practice led to decreases in brain activity in the bilateral parietal and premotor cortices. In contrast, no statistically significant changes were observed in primary sensorimotor areas. In secondary motor areas only, practice also resulted in decreased similarity of activation patterns between trained and untrained sequences. The similarity of the activation patterns among the trained sequences did not change. The preregistered predictions of practice-related changes in the variability of activation patterns across trials and in the estimates of brain structure were not supported by the data.

It is surprising that we were unable to detect training-related changes in the three structural measures we derived from the T1w scans, given that our experiment had more within-subject scans than most previous studies in the field and that the number of participants per group was on the upper end of the sample size range compared to similar studies (Draganski et al. 2004, 2006; Mårtensson et al. 2012; Lövdén et al. 2013). In addition, the reliabilities of our CT and GMV estimates were above moderate in the areas of interest. At present, the origin of learning-induced gray matter changes measured with MRI is still uncertain. An animal study which investigated the relationship between several neuron morphology metrics and the VBM signal found only a significant but weak association with spine density (Keifer et al. 2015). Another study of the effects of monocular deprivation in rats suggests that experience-dependent changes in GMV estimated with MRI mainly are the results of swelling astrocytes (Schmidt et al. 2021). It remains unknown whether these findings can be translated to humans learning a motor task. Our own post hoc simulations (Supplementary Information) of the relationship between statistical power and relative volumetric change suggest that, unless the relative changes are very large and possibly happening in several cellular constituents, they would be unlikely to be detected by vertex- or voxel-wise analyses (e.g., synaptic changes should be very extensive to trigger measurable macroscopic changes on their own). On the other hand, extant evidence indicates that vascular changes can induce changes in morphometric measures. For instance, it has been shown that a single-dose pharmacological manipulation that decreases cerebral blood flow in a localized manner alters VBM estimates in overlapping regions (Franklin et al. 2013; Ge et al. 2017). Other recent studies show task-related effects on MPRAGE images in humans (Månsson et al., 2018; Olivo et al., 2021). An intriguing explanation for the lack of structural alterations in the present study has to do with the structural sequence we used (MP2RAGE), which differs in a crucial way from the MPRAGE sequences that have typically been used in prior studies of experience-dependent plasticity in humans. Specifically, in MPRAGE sequences the T2* effects are present, whereas in flat images derived from MP2RAGE sequences T2* effects are mostly cancelled, by virtue of the division of the two volumes involved in order to remove the intensity bias (Marques et al. 2010; Tanner et al. 2012). Thus, if the learning-related structural changes observed in similar experimental designs based on MPRAGE sequences were predominantly of vascular origin, they would have been mostly cancelled out had MP2RAGE been used instead. Remarkably, even though the MP2RAGE sequence has been available for over 10 years (Marques et al. 2010) and it is becoming the recommended sequence for gray-white matter segmentation (Droby et al. 2021; Oliveira et al. 2021), no published studies have, to the best of our knowledge, shown longitudinal training-related structural changes on a similar time-scale using this MR sequence, which may be a symptom of its lower sensitivity to vascular alterations. Nevertheless, this interpretation remains speculative, and more research will be needed to elucidate this question.

Although executing the motor sequences triggered robust and widespread activations in visual, primary, and secondary motor cortices, and the nodes of the salience network, changes in activation were localized to bilateral parietal and premotor regions. Our finding of activation decreases in secondary cortical motor areas and absence of changes in the primary sensorimotor cortices agrees with recent inquiries (Wiestler and Diedrichsen 2013; Berlot et al. 2020), contradicting a number of older studies that reported increases (Karni et al. 1995, 1998; Grafton et al. 2002; Penhune and Doyon 2002; Floyer-Lea 2005; Lehéricy et al. 2005) or non-monotonic changes (Xiong et al. 2009; Ma et al. 2010). We can obviously not exclude that practicing tasks other than ours leads to decreases also in primary regions. Importantly, the task used in the present study was more challenging than those in previous similar studies in terms of motor demands (i.e., execution with non-dominant hand, relatively novel configural responses). Nevertheless, we observed no changes in activity in the primary sensorimotor cortices. In addition, in our study, subjects had no exposure to the motor sequences of the experiment prior to baseline imaging. To ensure that participants understood the mechanics of the task and were familiar with the process, they tested it only once during the information session several days prior to the baseline session, but the demonstration version of the task required executing easier sequences of 3 chords with transitions of only 1 or 2 finger changes at a time. Therefore, there was no meaningful pretraining of the actual experiment sequences that could have triggered changes before the first scanning session. We conclude that activity in secondary cortical motor areas declines from the beginning of the training period.

The set of regions in which activation exhibited changes for trained sequences corresponds to the dorsal attention network (DAN), or dorsal frontoparietal network (dFPN), which consists of the intraparietal sulcus and the frontal eye fields (Corbetta and Shulman 2002). These regions are activated simultaneously in a wide range of tasks, both motor (reaching, grasping, saccade production) and purely cognitive (spatial attention, mental rotation, working memory). Previous studies of finger sequence production have demonstrated that these regions encode sequences and sequence chunks (Wiestler and Diedrichsen 2013; Yokoi et al. 2018; Yokoi 2019; Berlot et al. 2020), and our multivariate analysis of the activity patterns also indicates that different sequences could be classified above chance level in these regions. Recent reviews have proposed that the overarching role of the DAN is the top-down modulation of attention (Vossel et al. 2014) or the emulation of actions (Ptak et al. 2017). One explanation for the decreases found in the DAN nodes is that, with practice, these areas become more efficient and therefore the demand for oxygen sinks (Berlot et al. 2020). A somewhat different perspective would be that, for sequences that have been intensively trained and memorized, the need to rely on spatial attention to read the sequences, subserved by the DAN, is much diminished. In the latter view, the changes do not reflect localized plastic change per se but the reduced need for involvement of this domain-general system after transitioning from controlled to automatic execution (Chein and Schneider 2012). Plastic change would happen at the level of the mechanisms selecting these regions in the early phases of skill learning.

The use of a control group, which former similar studies did not incorporate (Xiong et al. 2009; Ma et al. 2010; Wiestler and Diedrichsen 2013; Berlot et al. 2020), allowed us to establish that untrained sequences in the intervention group did not elicit significantly larger behavioral improvements or differential activity reductions with respect to the control group. This speaks to a lack of (measurable) generalization (i.e. the ability to transfer learned improvements to novel sequences (Krakauer et al. 2019)) from learning the trained sequences to the untrained ones in terms of both performance and functional activity in this task. However, the initial large changes occurring between the first and second sessions are an exception to this lack of generalization. It is likely that these initial changes relate to general aspects of getting acquainted with the task, such as selecting goals, developing online planning, or forming a strategy to perform the task (Krakauer et al. 2019; Spampinato and Celnik 2020; Ariani et al. 2021, 2022).

We found no evidence for the decreases in variability of activation patterns across trials predicted by the ESR model. A critical difficulty when comparing trial-wise variability of trained and untrained sequences is that, due to the low SNR that is typical of fMRI measurements, trial-wise pattern variability will have a large contribution from non-neural noise that cannot be disentangled from neural variability. For this reason, sensitivity to detect differences between variability of activation patterns will tend to be low. Moreover, an assumption that trained and untrained sequences have equivalent levels of non-neural noise must be made, which may not hold if systematic trained-untrained differences in non-neural signal are present.

Even though neural representations for different sequences were measurably distinct (i.e., classification accuracy was above chance) even when the sequences were not trained, practice did not lead to appreciable changes over time in cross-nobis dissimilarities between different (trained) sequences, as would be expected if the patterns became more distinct following training. Overall, our results are commensurate with the analysis of activation patterns for paced sequences by Berlot and colleagues (Berlot et al. 2020); they could not observe training-related effects on dissimilarity between patterns of different sequences that were executed at paced tempo either, only general increases in dissimilarity between trained- and untrained-sequence patterns in secondary motor regions.

We cannot rule out that the reason for the lack of structural findings is that the duration of the experiment was not long enough or that the task was not sufficiently demanding to elicit structural changes, compared to previous work. While some past studies have used periods of the order of months (Draganski et al. 2004; Matuszewski et al. 2021), the duration of our experiment was based on a former study of our group in which we were able to detect localized changes in motor areas (Wenger et al. 2017), together with a pilot study of the motor task that showed that the practice period extends beyond the point where performance reaches a plateau. We should therefore have been able to capture the whole process of exploration, selection, and refinement that is predicted by the ESR model. Longer periods of training (months or years) are possibly required to trigger measurable structural changes that give rise to differences between skilled and naïve groups (e.g., musicians vs. non-musicians; Schlaug et al. 2001; Gaser and Schlaug 2003). Alternatively, such differences may to some extent reflect selection effects (niche picking), with differences existing before practice. It is also possible that adaptive training procedures are required, but because our primary interest was to understand the dynamics of neural changes, we expressly avoided using an adaptive task that would have confounded the time-course of neural alterations.

We also note that other analysis approaches may be more effective in finding changes in neural representations. To obtain the activation patterns, we fitted a GLM to the functional data to derive one activation pattern per trial. This approach is bound to lose important timing information. A spatio-temporal approach that may potentially capture more nuanced aspects of neural representations would be considerably more complex to implement and remains as an avenue for future work. Lastly, in our study we opted for the use of a passive control group, which offers less control for unspecific aspects of learning the task than an active control group. However, we deem it unlikely that placebo or motivation effects can explain that the changes observed were localized only to motor-related areas.

In conclusion, training a paced configural sequence task with the non-dominant hand during a period of 6 weeks resulted in reduced activity in the DAN for the sequences that had been practiced, but neither in detectable activation changes in primary sensorimotor cortices nor in morphological changes. Practice also resulted in decreased similarity between the neural activation patterns of trained and untrained sequences in secondary, but not primary, motor areas.

## Disclosure of Competing Interests

The authors declare no competing interests.

## Acknowledgements

We would like to thank Marie Helsing for assistance with administrative matters. Sandra Persson, Agnes Wiberg, Theresa Ruwe, Ana Bernardi, Tobias Rosholm, Andrea Fingerhut, Tina Rastegar, Karen Kuckelkorn, William Fredborg, Danielle van Westen, Boel Hansson, Linda Wennberg, Johan Mårtensson and Karin Markenroth-Bloch played essential roles in recruitment, data collection, and data quality control. We thank Wietske van der Zwaag for kindly providing the MATLAB script to calculate the T1 relaxation times. We are grateful to Peter Mannfolk, Rita Almeida, Jonna Nilsson, Fredrik Ödegaard, Elisa Yaquian, and Eva Berlot for useful discussions. Philips Clinical Science provided tooling and support. Finally, we thank the participants for their commitment to the study.

## Funding

This work was supported by the Swedish Research Council [grant number 2018-01047] to M. L. and [grant number 2015-01717] to C. B., the Agence Nationale de la Recherche [grant number ANR-JC ANR-16-CE28-0008-01] to C. B. and the James McDonnell Foundation Award ‘Understanding human cognition’ [grant number 2021-3101] to C. B.

## Notes

### Competing Interest Statement

The authors have declared no competing interest.

## References

Amunts K, Schlaug G, Ja L, Steinmetz H, Schleicher A, Dabringhaus A, Zilles K. 1997. Motor Cortex and Hand Motor Skills: Structural Compliance in the Human Brain. Hum Brain Mapp. 5:206–215.

Ariani G, Kordjazi N, Pruszynski JA. 2021. The Planning Horizon for Movement Sequences. eNeuro. 8 (2):1–14.

Ariani G, Pruszynski JA, Diedrichsen J. 2022. Motor planning brings human primary somatosensory cortex into action-specific preparatory states. Elife. 11:1–20.

Avants BB, Epstein CL, Grossman M, Gee JC. 2008. Symmetric diffeomorphic image registration with cross-correlation: Evaluating automated labeling of elderly and neurodegenerative brain. Med Image Anal. 12:26–41.

Bengtsson SL, Nagy Z, Skare S, Forsman L, Forssberg H, Ullén F. 2005. Extensive piano practicing has regionally specific effects on white matter development. Nat Neurosci. 8:1148–1150.

Berlot E, Popp NJ, Diedrichsen J. 2020. A critical re-evaluation of fMRI signatures of motor sequence learning. E-life. 9:e55241.

Beukema P, Diedrichsen J, Verstynen T. 2019. Binding during sequence learning does not alter cortical representations of individual actions. J Neurosci. 39:6968–6977.

Changeux J-P. 1989. Neuronal models of cognitive functions. Cognition. 33:63–109.

Chein J, Schneider W. 2012. The Brain’s Learning and Control Architecture. Curr Dir Psychol Sci. 21:78–84.

Corbetta M, Shulman GL. 2002. Control of goal-directed and stimulus-driven attention in the brain. Nat Rev Neurosci. 3:201–215.

Dayan E, Cohen LG. 2011. Neuroplasticity subserving motor skill learning. Neuron. 72:443–454.

de Lange AMG, Bråthen Acs, Rohani DA, Grydeland H, Fjell AM, Walhovd KB. 2017. The effects of memory training on behavioral and microstructural plasticity in young and older adults. Hum Brain Mapp. 38:5666–5680.

de Manzano Ö, Ullén F. 2018. Same Genes, different brains: Neuroanatomical differences between monozygotic twins discordant for musical training. Cereb Cortex. 28:387–394.

Diedrichsen J, Ariani G, Berlot E. 2021. Estimating correlations between noisy activity patterns. A tricky problem with a generative solution. [WWW Document]. http://www.diedrichsenlab.org/BrainDataScience/noisy_correlation/index.htm.

Diedrichsen J, Kornysheva K. 2015. Motor skill learning between selection and execution. Trends Cogn Sci. 19:227–233.

Diedrichsen J, Kriegeskorte N. 2017. Representational models: A common framework for understanding encoding, pattern-component, and representational-similarity analysis, PLoS Computational Biology.

Diedrichsen J, Yokoi A, Arbuckle SA. 2017. Pattern component modeling: A flexible approach for understanding the representational structure of brain activity patterns. Neuroimage. 180:119–133.

Draganski B, Gaser C, Busch V, Schuierer G, Bogdahn U, May A. 2004. Changes in grey matter induced by training. Nature. 427:312–312.

Draganski B, Gaser C, Kempermann G, Kuhn HG, Bu C. 2006. Temporal and Spatial Dynamics of Brain Structure Changes during Extensive Learning. 26:6314–6317.

Droby A, Thaler A, Giladi N, Hutchison RM, Mirelman A, Bashat D Ben, Artzi M. 2021. Whole brain and deep gray matter structure segmentation: Quantitative comparison between MPRAGE and MP2RAGE sequences. PLoS One. 16:e0254597.

Edelman GM. 1987. Neural Darwinism: The theory of neuronal group selection. New York, NY, US: Basic books.

Esteban O, Markiewicz CJ, Blair RW, Moodie CA, Isik AI, Erramuzpe A, Kent JD, Goncalves M, Dupre E, Snyder M, Oya H, Ghosh SS, Wright J, Durnez J, Poldrack RA, Gorgolewski KJ. 2019. fMRIPrep: a robust preprocessing pipeline for functional MRI. Nat Methods. 16:111–116.

Fields RD. 2015. A new mechanism of nervous system plasticity: Activity-dependent myelination. Nat Rev Neurosci. 16:756–767.

Floyer-Lea A. 2005. Distinguishable Brain Activation Networks for Short- and Long-Term Motor Skill Learning. J Neurophysiol. 94:512–518.

Fonov V, Evans AC, Botteron K, Almli CR, Mckinstry RC. 2012. Unbiased Average Age-Appropriate Atlases for Pediatric Studies. Neuroimage. 54:313–327.

Franklin TR, Wang Z, Shin J, Jagannathan K, Suh JJ, Detre JA, O’Brien CP, Childress AR. 2013. A VBM study demonstrating “apparent” effects of a single dose of medication on T1-weighted MRIs. Brain Struct Funct. 218:97–104.

Fujimoto K, Polimeni JR, Kouwe AJW Van Der, Reuter M, Kober T, Benner T, Fischl B, Wald LL. 2014. Quantitative comparison of cortical surface reconstructions from MP2RAGE and multi-echo MPRAGE data at 3 and 7 T. Neuroimage. 90:60–73.

Gaser C, Schlaug G. 2003. Brain Structures Differ between Musicians and Non-Musicians. J Neurosci. 23:9240–9245.

Ge Q, Peng W, Zhang J, Weng X, Zhang Y, Liu T, Zang F, Wang Z. 2017. Short-term apparent brain tissue changes are contributed by cerebral blood flow alterations. PLoS One. 12:1–12.

Glasser MF, Coalson TS, Robinson EC, Hacker CD, Harwell J, Yacoub E, Ugurbil K, Andersson J, Beckmann CF, Jenkinson M, Smith SM, Van Essen DC. 2016. A multi-modal parcellation of human cerebral cortex. Nature. 536:171–178.

Gorgolewski K, Burns CD, Madison C, Clark D, Halchenko YO, Waskom ML, Ghosh SS. 2011. Nipype: A Flexible, Lightweight and Extensible Neuroimaging Data Processing Framework in Python. Front Neuroinform. 5:1–15.

Grafton ST, Hazeltine E, Ivry R. 1995. Functional Mapping of Sequence Learning in Normal Humans. J Cogn Neurosci. 7:497–510.

Grafton ST, Hazeltine E, Ivry RB. 2002. Motor sequence learning with the nondominant left hand: A PET functional imaging study. Exp Brain Res. 146:369–378.

Huang Y, Zhen Z, Song Y, Zhu Q, Wang S, Liu J. 2013. Motor Training Increases the Stability of Activation Patterns in the Primary Motor Cortex. PLoS One. 8.

Jenkinson M. 2003. Fast, Automated, N-Dimensional Phase-Unwrapping Algorithm. Magn Reson Med. 49:193–197.

Jenkinson M, Bannister P, Brady M, Smith S. 2002. Improved Optimization for the Robust and Accurate Linear Registration and Motion Correction of Brain Images. Neuroimage. 17:825–841.

Jensen SK, Yong VW. 2016. Activity-Dependent and Experience-Driven Myelination Provide New Directions for the Management of Multiple Sclerosis. Trends Neurosci. 39:356–365.

Karni A, Meyer G, Jezzard P, Adams MM, Turner R, Ungerleider LG. 1995. Functional MRI evidence for adult motor cortex plasticity during motor skill learning. Nature. 377:155–158.

Karni A, Meyer G, Rey-Hipolito C, Jezzard P, Adams MM, Turner R, Ungerleider LG. 1998. The acquisition of skilled motor performance: Fast and slow experience-driven changes in primary motor cortex. Proc Natl Acad Sci. 95:861–868.

Keifer OP, Hurt RC, Gutman DA, Keilholz SD, Gourley SL, Ressler KJ. 2015. Voxel-based morphometry predicts shifts in dendritic spine density and morphology with auditory fear conditioning. Nat Commun. 6:1–12.

Kilgard MP. 2012. Harnessing plasticity to understand learning and treat disease. Trends Neurosci. 35:715–722.

Klein A, Ghosh SS, Bao FS, Giard J, Stavsky E, Lee N, Rossa B, Reuter M, Neto EC. 2017. Mindboggling morphometry of human brains. PLOS Comput Biol. 13:e1005350.

Koo TK, Li MY. 2016. A Guideline of Selecting and Reporting Intraclass Correlation Coefficients for Reliability Research. J Chiropr Med. 15:155–163.

Krakauer JW, Hadjiosif AM, Xu J, Wong AL, Haith AM. 2019. Motor Learning. Compr Physiol. 9:613–663.

Ledoit O, Wolf M. 2004. A well-conditioned estimator for large-dimensional covariance matrices. 88:365–411.

Lehéricy S, Benali H, Van de Moortele P-F, Pélégrini-Issac M, Waechter T, Ugurbil K, Doyon J. 2005. Distinct basal ganglia territories are engaged in early and advanced motor sequence learning. Proc Natl Acad Sci. 102:12566–12571.

Lindenberger U, Lövdén M. 2019. Brain Plasticity in Human Lifespan Development: The Exploration – Selection – Refinement Model. Annu Rev Dev Psychol. 1:197–222.

Lövdén M, Garzón B, Lindenberger U. 2020. Human Skill Learning: Expansion, Exploration, Selection, and Refinement. Curr Opin Behav Sci. 36:163–168.

Lövdén M, Wenger E, Mårtensson J, Lindenberger U, Bäckman L. 2013. Structural brain plasticity in adult learning and development. Neurosci Biobehav Rev. 37:2296–2310.

Ma L, Wang B, Narayana S, Hazeltine E, Chen X, Robin DA, Fox PT, Xiong J. 2010. Changes in regional activity are accompanied with changes in inter-regional connectivity during 4 weeks motor learning. Brain Res. 1318:64–76.

Maguire EA, Gadian DG, Johnsrude IS, Good CD, Ashburner J, Frackowiak RSJ, Frith CD. 2000. Navigation-related structural change in the hippocampi of taxi drivers. PNAS. 97:4398–4403.

Makino H, Hwang EJ, Hedrick NG, Komiyama T. 2016. Circuit Mechanisms of Sensorimotor Learning. Neuron. 92:705–721.

Månsson KNT, Cortes DS, Manzouri A, Li T, Hau S, Fischer H. 2018. Viewing Pictures Triggers Rapid Morphological Enlargement in the Human Visual Cortex. 1–7.

Marques JP, Kober T, Krueger G, Zwaag W Van Der. 2010. MP2RAGE, a self bias-field corrected sequence for improved segmentation and T 1-mapping at high field. Neuroimage. 49:1271–1281.

Mårtensson J, Eriksson J, Bodammer NC, Lindgren M, Johansson M, Nyberg L, Lövdén M. 2012. Growth of language-related brain areas after foreign language learning. Neuroimage. 63:240–244.

Matuszewski J, Kossowski B, Bola Ł, Banaszkiewicz A, Gyger L, Kherif F, Szwed M, Frackowiak RS, Jednoróg K, Draganski B, Marchewka A. 2021. Brain plasticity dynamics during tactile Braille learning in sighted subjectslll: Multi-contrast MRI approach. Neuroimage. 227:117613.

Molina-Luna K, Hertler B, Buitrago MM, Luft AR. 2008. Motor learning transiently changes cortical somatotopy. Neuroimage. 40:1748–1754.

Mumford JA, Turner BO, Ashby FG, Poldrack RA. 2012. Deconvolving BOLD activation in event-related designs for multivoxel pattern classification analyses. Neuroimage. 59:2636–2643.

Nili H, Walther A, Alink A, Kriegeskorte N. 2020. Inferring exemplar discriminability in brain representations. PLoS One. 15:1–28.

Oliveira ÍAF, Roos T, Dumoulin SO, Siero JCW, Van W. 2021. Can 7T MPRAGE match MP2RAGE for gray-white matter contrast? Neuroimage. 240:118384.

Penhune VB, Doyon J. 2002. Dynamic Cortical and Subcortical Networks in Learning and Delayed Recall of Timed Motor Sequences. J Neurosci. 22:1397–1406.

Power JD, Barnes K, Snyder A. 2012. Spurious but systematic correlations in resting state functional connectivity MRI arise from head motion. Neuroimage. 59:2142–2154.

Ptak R, Schnider A, Fellrath J. 2017. The Dorsal Frontoparietal Network: A Core System for Emulated Action. Trends Cogn Sci. 21:589–599.

Reed A, Riley J, Carraway R, Carrasco A, Perez C, Jakkamsetti V, Kilgard MP. 2011. Cortical Map Plasticity Improves Learning but Is Not Necessary for Improved Performance. Neuron. 70:121–131.

Reuter M, Schmansky NJ, Rosas HD, Fischl B. 2012. Within-subject template estimation for unbiased longitudinal image analysis. Neuroimage. 61:1402–1418.

Reuter M, Tisdall MD, Qureshi A, Buckner RL, Kouwe AJW Van Der, Fischl B. 2015. Head motion during MRI acquisition reduces gray matter volume and thickness estimates. Neuroimage. 107:107–115.

Sampaio-Baptista C, Khrapitchev AA, Foxley S, Schlagheck T, Scholz J, Jbabdi S, Deluca GC, Miller KL, Taylor A, Thomas N, Kleim J, Sibson NR, Bannerman D, Johansen-berg H. 2013. Motor Skill Learning Induces Changes in White Matter Microstructure and Myelination. J Neurosci. 33:19499–19503.

Schlaug G, Israel B, Medical D. 2001. The Brain of Musicians. A Model for Functional and Structural Adaptation. Ann N Y Acad Sci. 930:281–299.

Schmidt S, Gull S, Herrmann K, Boehme M, Irintchev A, Urbach A, Reichenbach JR, Klingner CM, Gaser C, Witte OW. 2021. Experience-dependent structural plasticity in the adult brain: How the learning brain grows. Neuroimage. 225:117502.

Scholz J, Klein MC, Behrens TEJ, Johansen-Berg H. 2009. Training induces changes in white-matter architecture. Nat Neurosci. 12:1370–1371.

Spampinato D, Celnik P. 2020. Multiple Motor Learning Processes in Humans: Defining Their Neurophysiological Bases. Neurosci. 1–22.

Tanner M, Gambarota G, Kober T, Krueger G, Marques P, Newbould R, Erritzoe D. 2012. Fluid and White Matter Suppression With the MP2RAGE Sequence. J Magn Reson Imaging. 35:1063–1070.

Tustison N, Avants B, Cook P, Zheng Y, Egan A, Yushkevich P, Gee J. 2010. N4ITK: Improved N3 Bias Correction Nicholas. IEEE Trans Med Imaging. 29:1310–1320.

Vossel S, Geng JJ, Fink GR. 2014. Dorsal and ventral attention systems: Distinct neural circuits but collaborative roles. Neuroscientist. 20:150–159.

Walther A, Nili H, Ejaz N, Alink A, Kriegeskorte N, Diedrichsen J. 2016. Reliability of dissimilarity measures for multi-voxel pattern analysis. Neuroimage. 137:188–200.

Wenger E, Kühn S, Verrel J, Mårtensson J, Bodammer NC, Lindenberger U, Lövdén M. 2017. Repeated Structural Imaging Reveals Nonlinear Progression of Experience-Dependent Volume Changes in Human Motor Cortex. Cereb Cortex. 27:2911–2925.

Wiestler T, Diedrichsen J. 2013. Skill learning strengthens cortical representations of motor sequences. Elife. 2013:1–20.

Wong AL, Krakauer JW. 2019. Why Are Sequence Representations in Primary Motor Cortex So Elusive? Neuron. 103:956–958.

Xiong J, Ma L, Wang B, Narayana S, Duff EP, Egan GF, Fox PT. 2009. Long-term motor training induced changes in regional cerebral blood flow in both task and resting states. Neuroimage. 45:75–82.

Xu T, Yu X, Perlik AJ, Tobin WF, Zweig JA, Tennant K, Jones T, Zuo Y. 2009. Rapid formation and selective stabilization of synapses for enduring motor memories. Nature. 462:915–919.

Yang G, Pan F, Gan WB. 2009. Stably maintained dendritic spines are associated with lifelong memories. Nature. 462:920–924.

Yokoi A. 2019. Neural Organization of Hierarchical Motor Sequence Representations in the Human Neocortex Article Neural Organization of Hierarchical Motor Sequence Representations in the Human Neocortex. Neuron. 103:1–13.

Yokoi A, Arbuckle SA, Diedrichsen J. 2018. The role of human primary motor cortex in the production of skilled finger sequences. J Neurosci. 2798–17.

Yousry TA, Schmid UD, Alkadhi H, Schmidt D, Peraud A, Buettner A, Winkler P. 1997. Localization of the motor hand area to a knob on the precentral gyrus A new landmark. 141–157.

Zatorre RJ, Fields RD, Johansen-Berg H. 2012. Plasticity in gray and white: neuroimaging changes in brain structure during learning. Nat Neurosci. 15:528–536.

Zhang Y, Brady M, Smith S. 2001. Segmentation of Brain MR Images Through a Hidden Markov Random Field Model and the Expectation-Maximization Algorithm. IEEE Trans Med Imaging. 20:45–57.

