## Supplementary Information for "Cortical changes during the learning of sequences of simultaneous finger presses"

*Generation of motor sequences for the experimental task.* We generated all possible sequences of 5 chords with a Hamming distance of 3 between each transition (i.e., 3 fingers had to change between 2 consecutive combinations) and containing no repeated chords. We then found all groupings of 2 sets that could be formed with the sequences produced before where one set had 3 sequences (which would become trained sequences) and the other set (from which untrained sequences would be sampled) had at least 6 sequences, with the condition that there were no common transitions between trained and untrained sequences, as we assumed that the core learning component in the task were transitions rather than chords. We thus generated all the possible groupings with 3 trained and at least 6 untrained sequences with no common transitions between trained and untrained sequences. In all these groupings of sequences there were between 1 and 3 chords that were not shared between trained and untrained sequences (given the conditions above, groupings with no common chords are not possible). We then selected the configurations with the maximum possible number of chords that were not shared between trained and untrained (3 configurations). Each of them had 3 trained sequences and 6 untrained sequences. Finally, among these configurations we selected the 2 for which the frequency distribution of the different fingers was most similar between trained and untrained sequences. These are the configurations A and B mentioned in the main text. Code for generating the sequences can be found in https://github.com/benjamingarzon/SeqLearn/blob/master/stats/SequenceStructure.Rmd.

*Consistency of MT curves.* MT time-courses were consistent within subjects (Figure 2 A), such that curves corresponding to pairs of sequences executed by the same subject were more similar than curves of pairs corresponding to different subjects (correlation between different learning curves, averaged across sequence pairs: within-subject mean correlation = 0.63 (SD = 0.22), between-subject mean correlation = 0.32 (SD = 0.27), Wilcoxon rank sum test W = 85524, p < 1e-10).

*Analysis of longitudinal relaxation (T1) times.* Approximate longitudinal relaxation values were obtained from the flat images using a protocol-specific lookup table relating pixel value and T1 values described in (Marques et al. 2010) and implemented in MATLAB. We assumed a constant B1 field, which is an acceptable assumption in the flat images within the cortical regions we probed because of the inherent B1-correction of the MP2RAGE sequence.

The T1 maps were sampled (mri_vol2surf Freesurfer tool) at a relative cortical depth (downward distance from the pial surface) corresponding to 20 %, 40 %, 60 % and 80 % of CT, yielding one T1 cortical map for each depth. Like the CT maps, the T1 cortical maps were warped onto the fsaverage surface and smoothed with a 10 mm FWHM kernel.

Reliability of T1 values, computed vertex-wise as ICC values in the same way as for CT, was poor to moderate (median = 0.5, SD = 0.13; Supplementary Figure 3 C, F), possibly reflecting residual B1+ inhomogeneity. Statistical tests were equivalent to those for CT and GMV, albeit adding relative cortical depth as a regressor. The univariate analysis did not identify any clusters displaying a pattern consistent with gradual differential increases, decreases or non-linear progression in the intervention group with respect to the control group in T1 relaxation time.

*Simulations of relative changes vs. power.* Based on the data we had collected, we simulated synthetic relative structural changes to estimate the relationship between power and the amount of structural changes, with the aim to ascertain whether the amount of growth of different cellular constituents necessary to generate detectable effects on morphometric measures is plausible. We created two synthetic subject groups by sampling 30 subjects with replacement from the acquired control subjects. For each of the selected subjects, we estimated mean and standard deviation of CT in 200 randomly selected vertices and used these estimates to generate normally distributed data in as many synthetic vertices and 7 different sessions. In one of the groups (synthetic intervention group), we added a relative change with either a linear, asymptotic or quadratic time-course, and used an LMM with the same model to estimate the p-value for a group x session interaction. We repeated the sampling procedure 500 times and derived an estimate of statistical power as the fraction of the samples in which the p-value was below 0.001 (this value would correspond to a lenient uncorrected level of significance, e.g., Wenger et al. 2021) and averaging this fraction over synthetic vertices. The level of relative change (peak of the time course) in tissue volume was varied between 0 and 5 % so that we could estimate its relationship with statistical power, and a cubic root correction was applied to translate it to relative change in CT (this assumes isotropic growth). The relative size of experience-dependent cortical morphometric changes lies below 5% (Draganski et al. 2004, 2006; Mårtensson et al. 2012; Wenger et al. 2016). For further comparison, annual percent reductions in CT estimated from a large dataset were around 1% during development and between 0.1% and 0.5% in early and middle adulthood (Fjell et al. 2015). Lastly, to take into account the fact that gray matter is not homogeneous and that changes should not be assumed to occur in the whole tissue volume, we divided the level of change applied by published estimates of the relative proportion of the major cellular constituents of gray matter, namely axonal collaterals (29%), dendrites (26%), soma (11%), astrocytes (9%), synapses (5.7%) and oligodendrocytes (2.5%) (Kassem et al. 2013), and rendered a curve of power vs. relative change for each of these components. Note that these estimates are derived from rodent data and only used here for illustrative purposes, but we doubt that they would be so different in humans so as to invalidate our main point. We repeated the analysis above also for GMV values (no correction was applied as these estimates already reflect volume), and for the case of having only 3 sessions (sessions 1, 4 and 7), since most similar studies included only 2 or 3 sessions. Additionally, we performed all the simulations above again but this time attempting to reproduce a ROI-based analysis. In this case we averaged the simulated structural measures for all the 200 vertices/voxels first and set the significance threshold to 0.05 when deriving an estimate of statistical power. This analysis was done assuming that either 33%, 66% or 100% of the ROI voxels/vertices presented the effect in order to evaluate the robustness of the results in different scenarios with a varying proportion of the ROI being affected by changes.

The results of the simulation show that for vortex-/vertex-wise analyses, if the relative size of the changes was around what typically has been reported (< 5%), the power to detect these changes would in general be quite low (Supplementary Figure 4 A, B), particularly for CT. Moreover, the relative growth for individual constituents resulting in those changes in total CT / GMV should be much larger, above 20%, to achieve desirable levels of statistical power given the inherent reliability of the data. Thus, these results support the view that the observed changes in morphometric estimates in vertex-/voxel-wise analyses are unlikely to be caused by the growth of any of these tissue compartments individually, as this would demand unrealistically large changes in them. Naturally, ROI analyses are more powerful (Supplementary Figure 4 C - H), albeit obviously at the expense of lower spatial specificity. Unless effects are present in a large proportion of the ROI (E – H), they still require large changes in individual cellular constituents to be detected.

### References

Draganski B, Gaser C, Busch V, Schuierer G, Bogdahn U, May A. 2004. Changes in grey matter induced by training. Nature. 427:312–312.

Draganski B, Gaser C, Kempermann G, Kuhn HG, Bu C. 2006. Temporal and Spatial Dynamics of Brain Structure Changes during Extensive Learning. 26:6314–6317.

Fjell AM, Grydeland H, Krogsrud SK, Amlien I, Rohani DA, Ferschmann L. 2015. Development and aging of cortical thickness correspond to genetic organization patterns. PNAS. 112:1–6.

Glasser MF, Coalson TS, Robinson EC, Hacker CD, Harwell J, Yacoub E, Ugurbil K, Andersson J, Beckmann CF, Jenkinson M, Smith SM, Van Essen DC. 2016. A multi-modal parcellation of human cerebral cortex. Nature. 536:171–178.

Kassem MS, Lagopoulos J, Stait-Gardner T, Price WS, Chohan TW, Arnold JC, Hatton SN, Bennett MR. 2013. Stress-induced grey matter loss determined by MRI is primarily due to loss of dendrites and their synapses. Mol Neurobiol. 47:645–661.

Marques JP, Kober T, Krueger G, Zwaag W Van Der. 2010. MP2RAGE, a self bias-field corrected sequence for improved segmentation and T 1 -mapping at high field. Neuroimage. 49:1271–1281.

Mårtensson J, Eriksson J, Bodammer NC, Lindgren M, Johansson M, Nyberg L, Lövdén M. 2012. Growth of language-related brain areas after foreign language learning. Neuroimage. 63:240–244.

Oldfield RC. 1971. The assessment and analysis of handedness: The Edinburgh inventory. Neuropsychologia. 9:97–113.

Wenger E, Kühn S, Verrel J, Mårtensson J, Bodammer NC, Lindenberger U, Lövdén M. 2016. Repeated Structural Imaging Reveals Nonlinear Progression of Experience-Dependent Volume Changes in Human Motor Cortex. Cereb Cortex. 1–15.

Wenger E, Papadaki E, Werner A, Kühn S, Lindenberger U. 2021. Observing Plasticity of the Auditory System : Volumetric Decreases Along with Increased Functional Connectivity in Aspiring Professional Musicians. 1–14.

### Supplementary Figures

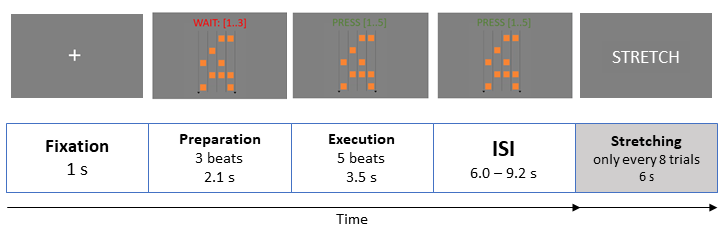

**Supplementary Figure 1. Timeline of an fMRI trial.** After a fixation cross was shown for 1 s, subjects were prompted to wait during 3 preparation beats that signaled the predefined tempo (0.7 s / beat) at which to press the keys. This preparation phase was followed by an execution phase of 5 beats in which subjects were supposed to press the sequence of chords at the shown tempo. The beats were indicated by a counter. Following the execution phase there was a pseudorandom exponentially distributed inter-trial-interval with mean 7.4 s and truncated between 6.0 s and 9.2 s, counterbalanced across subjects. To improve participants’ comfort during the task and avoid that superfluous movements produced undesired BOLD signal fluctuations, after every 8 trials subjects were given 6 s to stretch their hand, indicated by the text ‘STRETCH’ displayed on the screen.

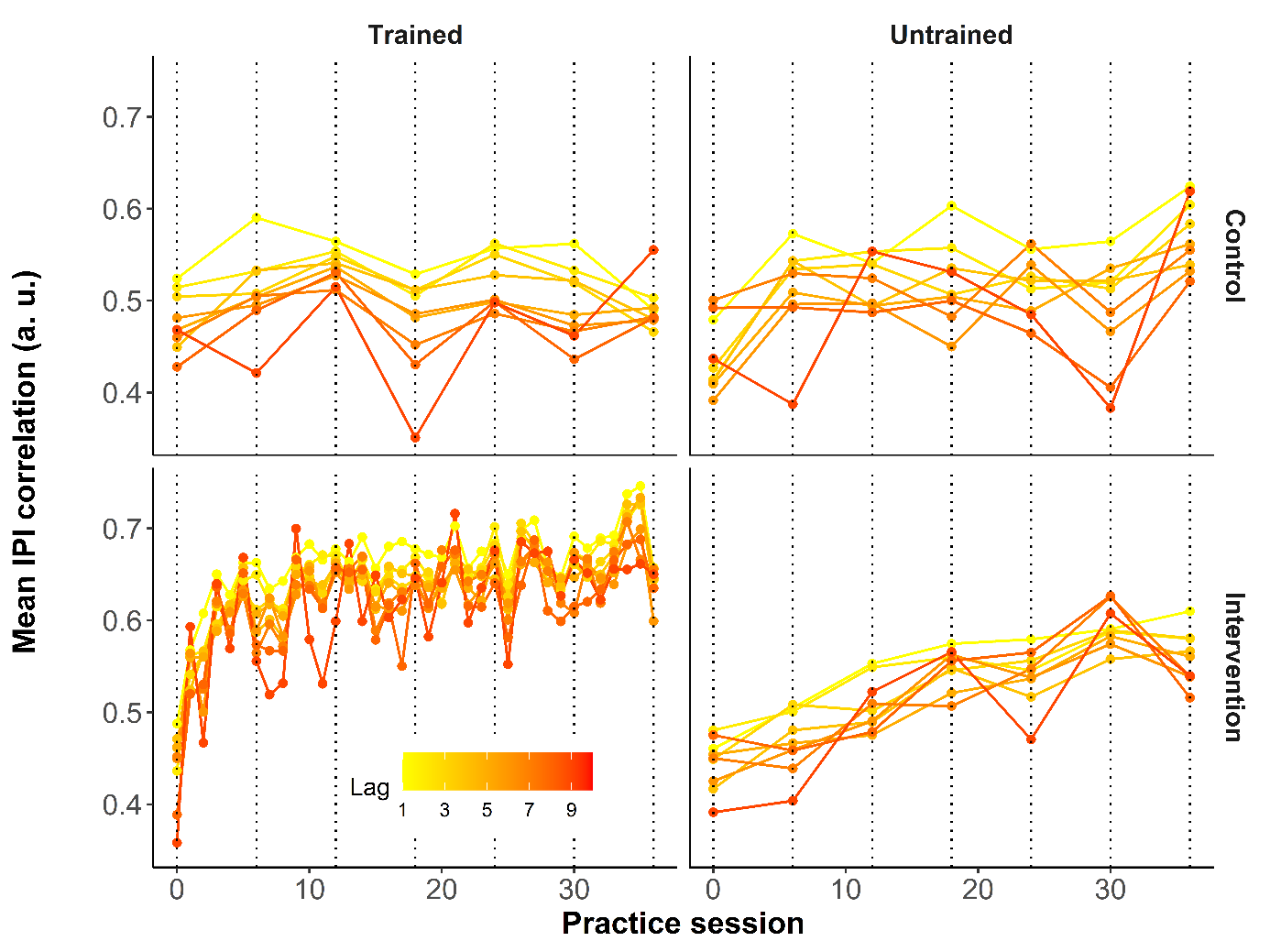

**Supplementary Figure 2. Correlation of inter-press intervals (IPIs).** Practice of the trained sequences resulted in increased correlation between the IPIs of different executions within the same session for those particular sequences. Correlation between IPIs for pairs of trials at different lags (each trace corresponds to a lag). The practice sessions are also shown, and the dotted vertical lines correspond to the 7 test sessions (cfr. Figure 2 in the main text).

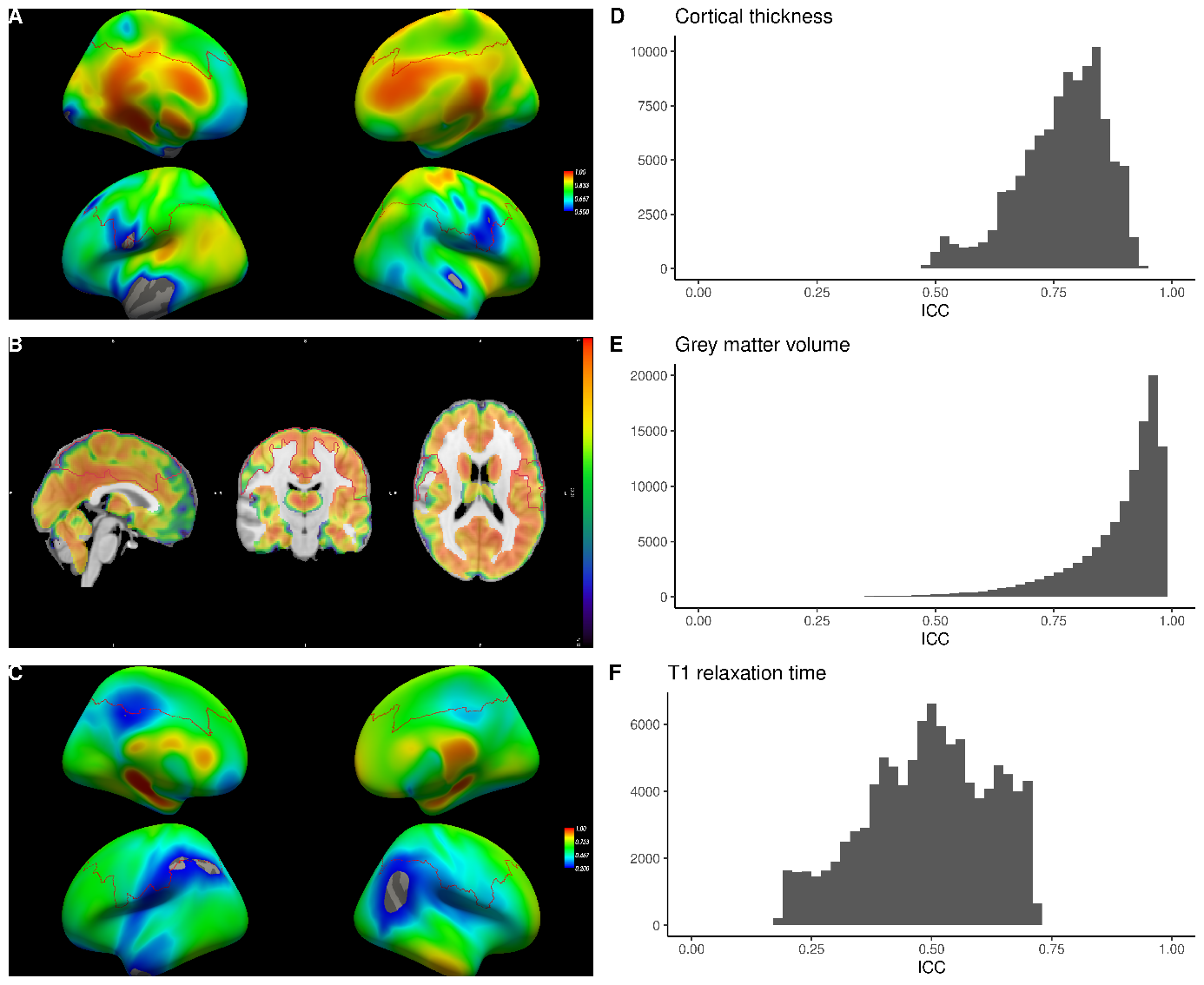

**Supplementary Figure 3. Reliability of structural measures.** Maps of intra-class correlation coefficients across vertices/voxels for cortical thickness (A), gray matter volume (B) and T1 relaxation time (C). The red outline depicts the ROI for the structural analyses reported in the main text (see also Supplementary Figure 8). The histograms show the distribution of ICC within this ROI for cortical thickness (D), gray matter volume (E) and T1 relaxation time (F). Note that the color scale in C is different from those in A and B due to the lower ICC values for T1 relaxation time. Analyses of cortical thickness and gray matter volume are reported in the main text and the analysis of T1 values is reported in the Supplementary Information.

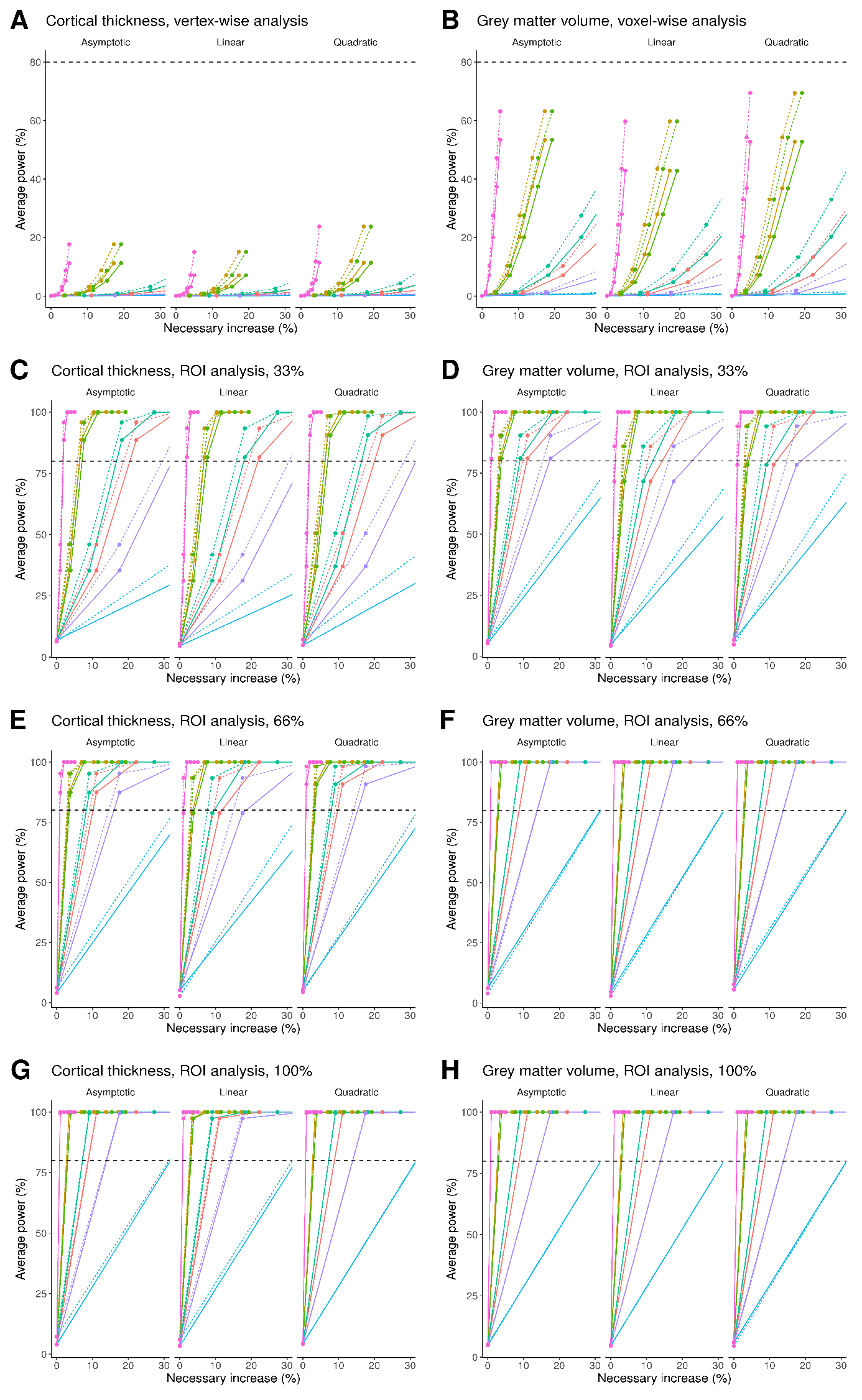

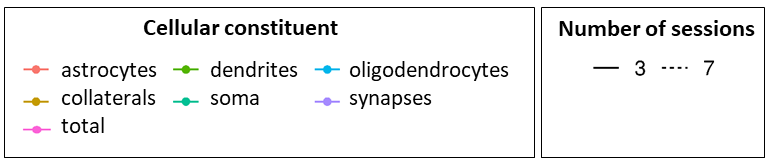

**Supplementary Figure 4. Relationship between statistical power to detect a significant group x session interaction in morphometric measures and synthetic relative changes.** Based on the acquired data for the control group, we simulated changes in a synthetic intervention group in cortical thickness (A) and gray matter volume (B) and at different relative levels, between 0 and 5% varying in steps of 1% (each dot). We fitted linear mixed models to each simulated instance and measured the power to detect such changes for an uncorrected significance level of α = 0.001 (see text). The procedure was performed assuming that changes could be linear, asymptotic or quadratic, and with 3 or 7 scanning sessions. Based on approximate proportions for different cellular constituents of gray matter, we estimated the corresponding curves (marked with different colors) to illustrate the relative morphometric changes needed to achieve a certain statistical power. Results of the simulations of equivalent ROI analyses, obtained by averaging the structural measures across vertices and setting α = 0.05, are displayed in the remaining panels (C, E, G for CT) and (D, F, H for GMV). In panels C, D the simulated effect was present in 33% of vertices/voxels, for E, F in 66% and for G, H in 100%. The horizontal dashed line in each panel indicates 80% power.

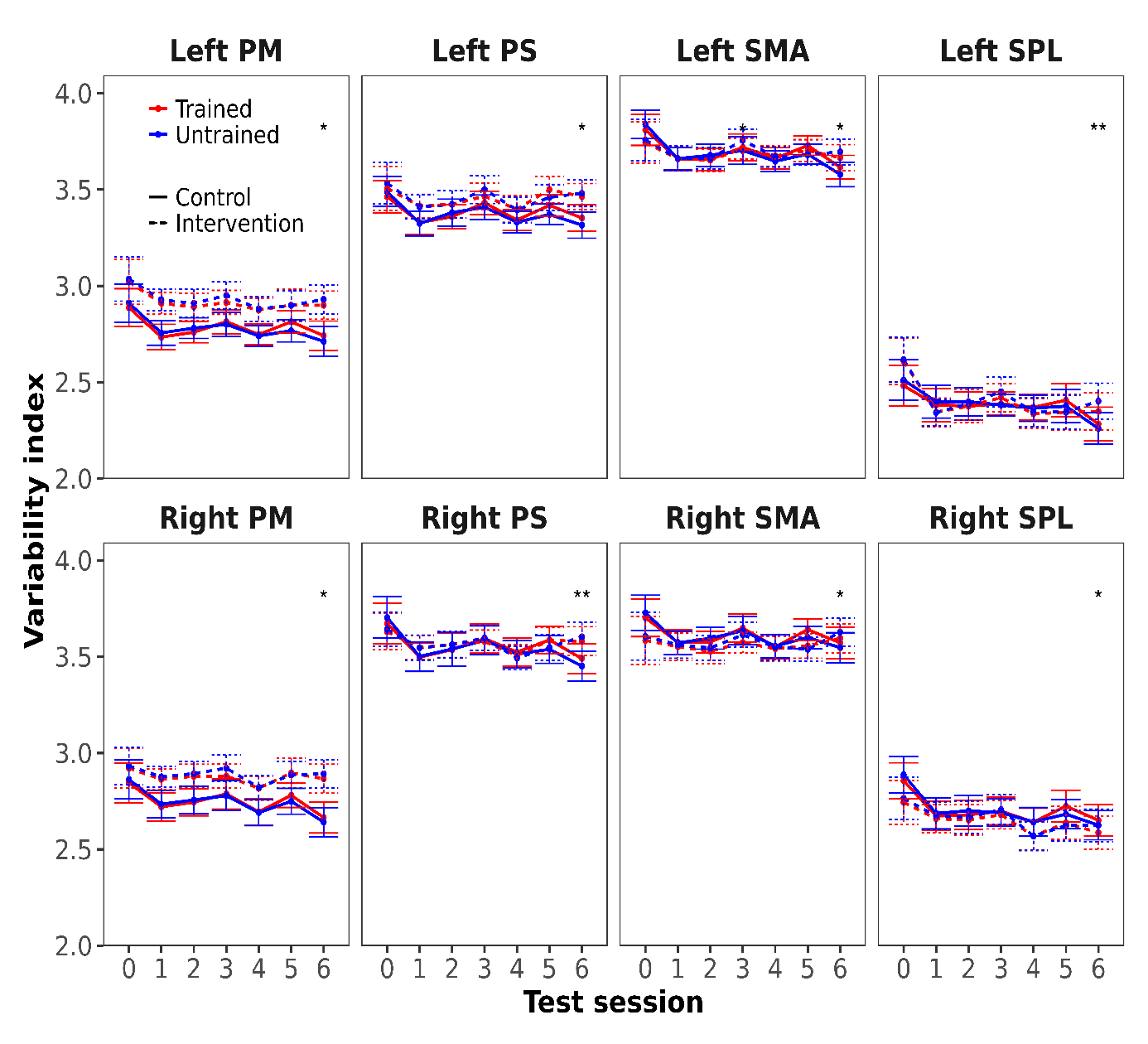

**Supplementary Figure 5. Neural pattern variability index over time in each of the tested ROIs.** No changes in variability between trained and untrained sequences could be detected. ROI: region-of-interest; PM: premotor; PS: primary sensorimotor; SMA: supplementary motor area; SPL: superior parietal lobule; ‘***’ significant FDR-corrected for ROIs and sessions, q< 0.05; ‘**’ significant FDR-corrected for sessions, q< 0.05 ‘*’ significant uncorrected, p < 0.05.

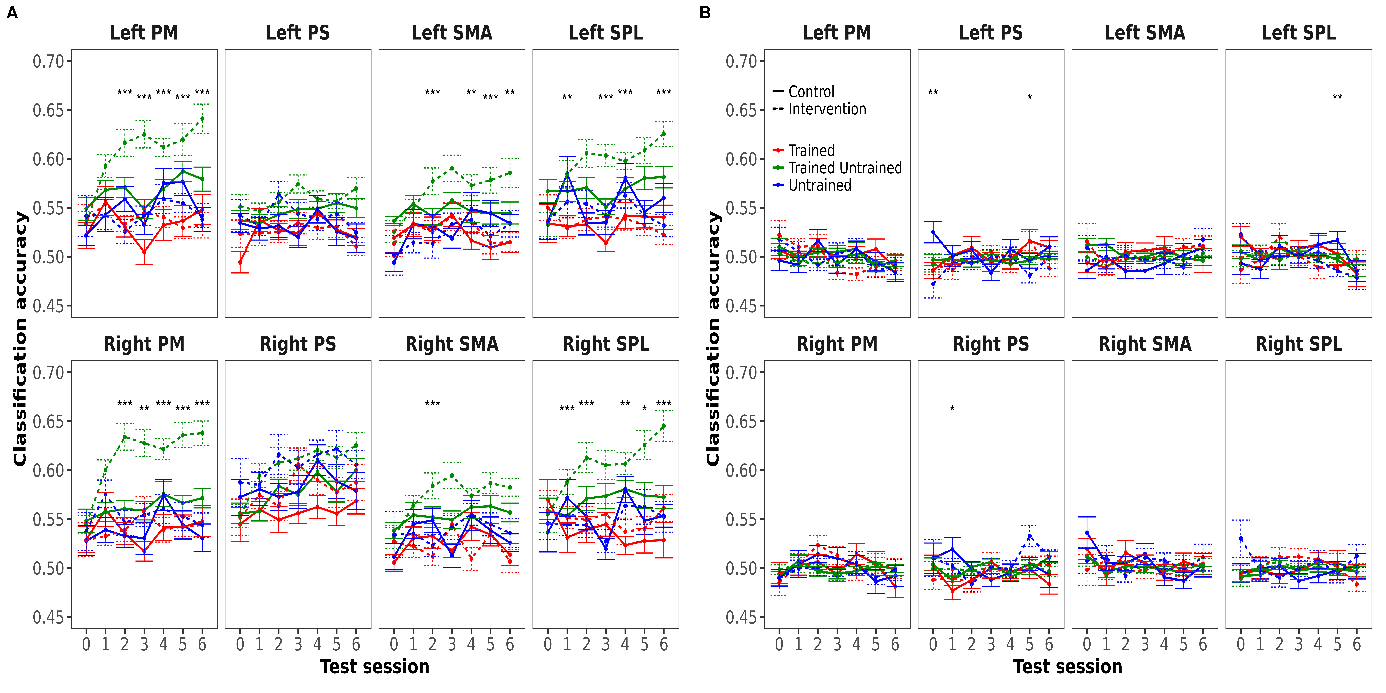

**Supplementary Figure 6. Cross-validated classification accuracy for a support vector machine classifier trained on pairs of neural patterns to discriminate between different sequences.** A) Accuracy was averaged separately for different categories of sequence pairs (i.e., both sequences in the pair were trained, both untrained or one was trained and the other was untrained). Asterisks indicate a significant interaction of group x practice (trained - untrained vs. untrained - untrained) in that session. B) Results of an equivalent analysis after shuffling the labels randomly. Accuracies dropped to chance level (0.5 probability) and there was no dependence on session for any of the sequence categories, indicating that accuracies above 0.5 and patterns of increase over time displayed in (A) could not arise randomly. PM: premotor; PS: primary sensorimotor; SMA: supplementary motor area; SPL: superior parietal lobule; ‘***’ significant FDR-corrected for ROIs and sessions, q< 0.05; ‘**’ significant FDR-corrected for sessions, q < 0.05 ‘*’ significant uncorrected, p < 0.05.

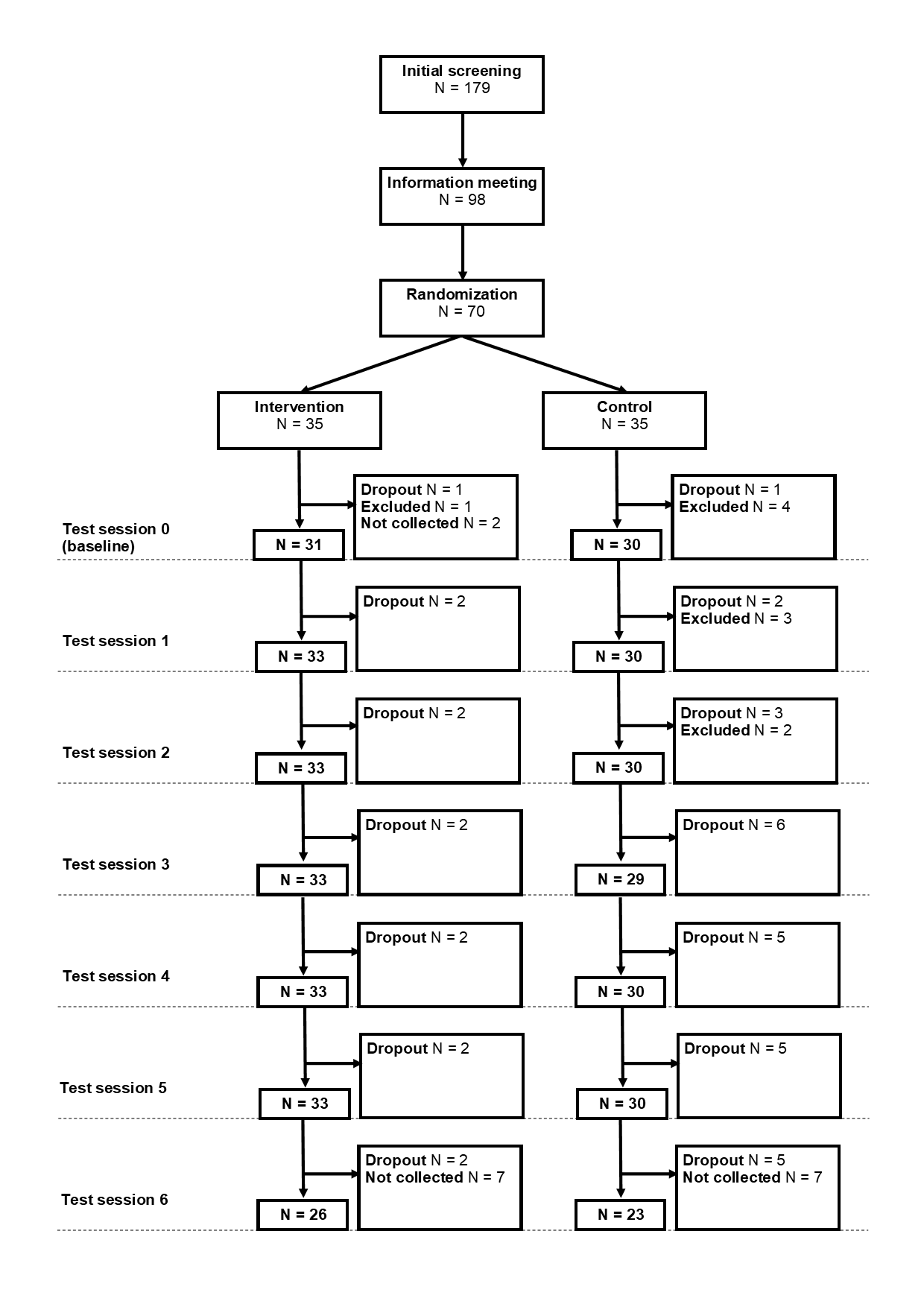

**Supplementary Figure 7. Recruitment, randomization and attrition.** The flow chart shows the number of subjects involved in the different study stages: subjects were initially screened via a telephone call and those fulfilling the study criteria invited to an information meeting. Seventy of those consenting to participate in the study were randomized in either an intervention or a control group. Subjects were excluded from the analysis if they attended less than 4 test sessions. Some measurements were not collected due to technical issues, as for instance in the last week of the 4^th^ wave of the study, which is reflected in the lower numbers for the last test session.

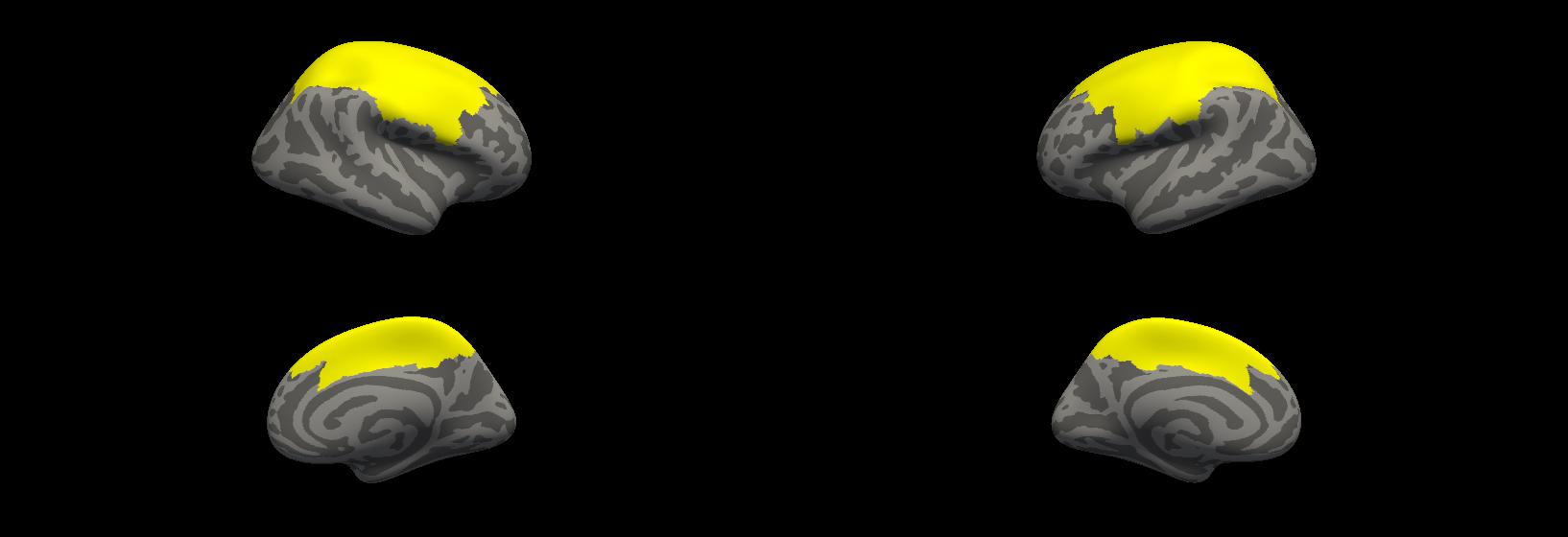

**Supplementary Figure 8. Mask used to constrain the structural analyses.** The parcels from the Human Connectome Project’s Multi-modal Cortical Parcellation (Glasser et al. 2016) that were combined to form the mask are shown in Supplementary Table 4.

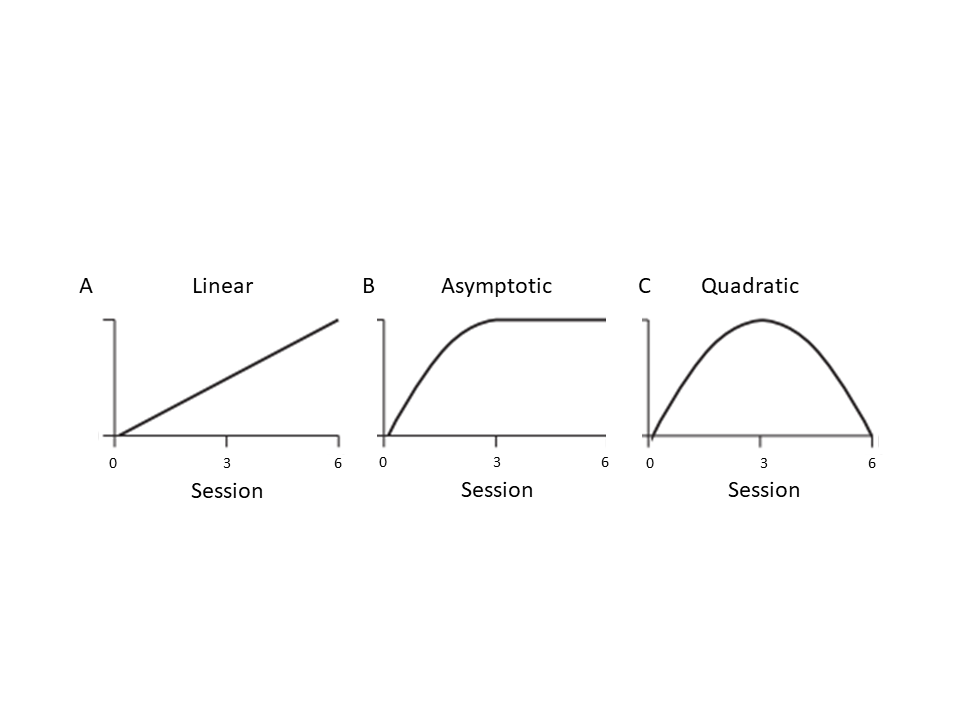

**Supplementary Figure 9. Models of structural and functional change.** Schematic representation of the linear (A), asymptotic (B) and quadratic (C) models that were tested.

### Supplementary Tables

**Supplementary Table 1. Inclusion and exclusion criteria.**

**Inclusion criteria:**

- Age between 20 and 30 years
- Availability to attend all MR sessions and practice the task at home during the experiment period
- Normal or corrected vision
- Fluent Swedish or English (required to understand the instructions)
- Enough mobility to lie flat and still in the MRI scanner for up to one hour
- Right-handedness (minimum score of 8 in the modified version of the Edinburgh Handedness Inventory; Oldfield 1971)
- Weight < 120 kg

**Exclusion criteria:**

- Having motor impairments in the left hand, fingers or arm (e.g., chronic pain, arthritis)
- Having played musical instruments or performed any other activity which involved fine motor skills with the left hand (magic tricks, video games) at some point in life for more than 12 consecutive months
- Having played musical instruments or performed frequently any other activity involving fine motor skills with the left hand in the past 5 years
- Having participated in other research studies involving fine motor skill training
- Neurological diseases, including Parkinson’s disease, dementia and epilepsy
- Previous or current cardiovascular diseases (except for hypertension)
- History of brain damage or stroke
- Type I or pharmacologically-treated type II diabetes (type II diabetes allowed if treated only with dietary intervention)
- Current cancer (allowed if more than one year has passed since treatment end)
- Psychiatric illness (history of mild to moderate depression and/or anxiety allowed)
- History of head trauma resulting in loss of consciousness
- ADHD/ADD
- Presence of metal in the body (including piercings or make-up that cannot be removed)
- Claustrophobia
- Sound sensitivity (due to MRI noise)
- Previous or ongoing cardiovascular disease (except hypertension)
- Being pregnant

| **Configuration** | **Trained** | **Untrained** |
| --- | --- | --- |
| **A** | "1 3 - 2 - 3 4 - 1 2 3 - 1 4"  "3 4 - 2 - 1 3 - 2 3 4 - 1 4"  "1 4 - 2 3 4 - 1 2 - 4 - 2 3" | "2 4 - 1 - 1 2 3 4 - 2 - 1 4"  "1 2 - 3 - 1 2 3 4 - 4 - 1 3"  "2 4 - 3 - 1 2 - 2 3 4 - 1 3"  "1 3 - 4 - 1 2 3 4 - 1 - 3 4"  "2 4 - 1 2 3 - 3 4 - 1 - 2 3"  "1 4 - 2 - 1 2 3 4 - 3 - 2 4" |
| **B** | "3 4 - 1 - 2 3 - 4 - 1 2 3 4"  "2 3 - 1 - 1 2 3 4 - 2 - 3 4"  "2 - 1 2 3 4 - 1 - 2 4 - 1 2 3" | "1 2 3 - 3 4 - 2 - 1 4 - 3"  "2 3 4 - 1 2 - 4 - 1 3 - 2"  "1 2 3 - 2 4 - 3 - 1 4 - 2 3 4"  "1 2 - 3 - 2 4 - 1 - 3 4"  "1 2 3 - 1 4 - 2 - 1 3 - 4"  "3 - 1 2 - 2 3 4 - 1 4 - 1 2 3" |

**Supplementary Table 2. Sequences used in the task.** There were two different configurations that were randomly assigned to participants within each experimental group (i.e., 17 subjects in each group received configuration A and the remaining 18 configuration B). Tests in the scanner involved only 2 trained and 2 untrained sequences, and the A and B subgroups were further split into 2 subgroups each (A1/A2/B1/B2), depending on which 2 of the 3 trained sequences the subjects were tested on. Each configuration consisted of 3 *trained* sequences (which subjects in the intervention group practiced at home) and 6 *untrained* sequences. In the table, a sequence, for example "1 3 - 2 - 3 4 - 1 2 3 - 1 4", is given by 5 chords separated by hyphens. A chord is indicated by a list of the left-hand fingers involved (1=pinky, 2=ring, 3=middle, 4=index). In this example sequence, for the first chord (“1 3”) subjects needed to press with the pinky and middle finger simultaneously. For the second one, “2”, they only had to press with the ring finger.

**Supplementary Table 3. Overview of characteristics of the task for each phase and session.** At home, the unpaced phase preceded the paced phase. At the MR facility, the unpaced phase (outside the scanner) preceded the paced phase (inside the scanner) in the baseline session, whereas in the remaining sessions phase order was swapped.

| **Session type** | **Location** | **Sessions (baseline = 0)** | **Subject group** | **Pace type** | **Number of trials per sequence** | **Feedback type** | **Number of different sequences** |
| --- | --- | --- | --- | --- | --- | --- | --- |
| **Training - unpaced phase** | Home | 1 - 5, 7 -11, 13 – 17, 19 – 23, 25 – 29, 31 - 35 | Intervention | Unpaced | 20 | Correct / incorrect, missed, too slow, score (execution speed) | 3 trained |
| **Training - paced phase** | Home | 1 - 5, 7 -11, 13 – 17, 19 – 23, 25 – 29, 31 - 35 | Intervention | Paced (screen counter + auditory beat) | 5 | Correct / incorrect, missed, too slow / too fast | 3 trained |
| **Testing – paced phase** | MR center, inside scanner | 0, 6, 12, 18, 24, 30, 36 | Intervention and control | Paced (screen counter) | 20 | None | 2 trained, 2 untrained |
| **Testing – unpaced phase** | MR center, outside scanner | 0, 6, 12, 18, 24, 30, 36 | Intervention and control | Unpaced | 20 | Correct / incorrect, missed, too slow, score (execution speed) | 2 trained, 3 untrained |

**Supplementary Table 4.** Parcels from the Human Connectome Project’s Multi-modal Cortical Parcellation (Glasser et al. 2016) that were combined to form the mask (ROI_surf_) used to constrain the structural analyses. This mask was subsequently projected onto MNI space to create a mask to constrain the volume-based analyses (ROI_vol_).

| **Left hemisphere** | **Right hemisphere** |
| --- | --- |
| L_4_ROI L_3b_ROI L_FEF_ROI L_PEF_ROI L_55b_ROI L_SFL_ROI L_PCV_ROI L_7Pm_ROI L_5m_ROI L_5mv_ROI L_23c_ROI L_5L_ROI L_24dd_ROI L_24dv_ROI L_7AL_ROI L_SCEF_ROI L_6ma_ROI L_7Am_ROI L_7PL_ROI L_7PC_ROI L_LIPv_ROI L_VIP_ROI L_MIP_ROI L_1_ROI L_2_ROI L_3a_ROI L_6d_ROI L_6mp_ROI L_6v_ROI L_p24pr_ROI L_a24pr_ROI L_8Av_ROI L_8Ad_ROI L_8BL_ROI L_8C_ROI L_IFJp_ROI L_LIPd_ROI L_6a_ROI L_i6-8_ROI L_s6-8_ROI L_PFt_ROI L_AIP_ROI L_IP2_ROI L_IP1_ROI L_PFop_ROI L_31a_ROI L_p32pr_ROI L_6r_ROI L_8BM_ROI | R_4_ROI R_3b_ROI R_FEF_ROI R_PEF_ROI R_55b_ROI R_SFL_ROI R_PCV_ROI R_7Pm_ROI R_5m_ROI R_5mv_ROI R_23c_ROI R_5L_ROI R_24dd_ROI R_24dv_ROI R_7AL_ROI R_SCEF_ROI R_6ma_ROI R_7Am_ROI R_7PL_ROI R_7PC_ROI R_LIPv_ROI R_VIP_ROI R_MIP_ROI R_1_ROI R_2_ROI R_3a_ROI R_6d_ROI R_6mp_ROI R_6v_ROI R_p24pr_ROI R_a24pr_ROI R_8Av_ROI R_8Ad_ROI R_8BL_ROI R_8C_ROI R_IFJp_ROI R_LIPd_ROI R_6a_ROI R_i6-8_ROI R_s6-8_ROI R_PFt_ROI R_AIP_ROI R_IP2_ROI R_IP1_ROI R_PFop_ROI R_31a_ROI R_p32pr_ROI R_6r_ROI R_8BM_ROI |

**Supplementary Table 5. Statistics for behavioral measures**. Estimates for the group (intervention/control) x practice (trained/untrained) interaction for each of the test sessions and the different behavioral measures, obtained by fitting linear mixed models (LMMs) with subjects as random effects. Note that for the analysis of the number of correct trials, we used a generalized LMM with binomial distribution, and the statistic returned by the fitting routine is normally distributed, as opposed to the other analyses, were statistics follow a Student’s t distribution.

| **Name** | **Session** | **Estimate** | **Std. Error** | **df** | **t** | **p(uncorrected)** | **p(corrected)** | **Significance** |
| --- | --- | --- | --- | --- | --- | --- | --- | --- |
| **Median MT** | 0 | 0.030 | 0.039 | 215.570 | 0.769 | 0.221 | 0.221 | n. s. |
|  | 6 | 0.091 | 0.029 | 232.854 | 3.085 | 0.001 | 0.003 | ** |
|  | 12 | 0.097 | 0.032 | 230.515 | 3.047 | 0.001 | 0.003 | ** |
|  | 18 | 0.082 | 0.033 | 219.248 | 2.490 | 0.006 | 0.011 | ** |
|  | 24 | 0.056 | 0.032 | 234.540 | 1.758 | 0.039 | 0.046 | ** |
|  | 30 | 0.111 | 0.030 | 231.549 | 3.675 | 1.19e-04 | 8.34e-04 | ** |
|  | 36 | 0.086 | 0.043 | 180.194 | 1.993 | 0.023 | 0.032 | ** |
| **MT standard deviation** | 0 | -0.114 | 0.105 | 218.026 | -1.086 | 0.861 | 0.861 | n. s. |
|  | 6 | 0.107 | 0.104 | 235.438 | 1.034 | 0.151 | 0.264 | n. s. |
|  | 12 | 0.284 | 0.107 | 233.184 | 2.658 | 0.004 | 0.028 | ** |
|  | 18 | -0.029 | 0.115 | 223.740 | -0.252 | 0.600 | 0.699 | n. s. |
|  | 24 | 0.242 | 0.116 | 236.750 | 2.083 | 0.019 | 0.065 | * |
|  | 30 | 0.189 | 0.119 | 231.380 | 1.589 | 0.056 | 0.131 | n. s. |
|  | 36 | 0.102 | 0.146 | 181.926 | 0.696 | 0.243 | 0.341 | n. s. |
| **IPI correlation** | 0 | 0.031 | 0.104 | 213.937 | 0.299 | 0.618 | 0.618 | n. s. |
|  | 6 | -0.288 | 0.107 | 234.093 | -2.697 | 0.003 | 0.006 | ** |
|  | 12 | -0.302 | 0.111 | 232.642 | -2.728 | 0.003 | 0.006 | ** |
|  | 18 | -0.345 | 0.117 | 227.862 | -2.944 | 0.002 | 0.006 | ** |
|  | 24 | -0.260 | 0.116 | 236.173 | -2.247 | 0.012 | 0.014 | ** |
|  | 30 | -0.283 | 0.113 | 232.825 | -2.508 | 0.006 | 0.008 | ** |
|  | 36 | -0.427 | 0.138 | 181.575 | -3.103 | 9.59e-04 | 0.006 | ** |
| **Name** | **Session** | **Estimate** | **Std. Error** | **df** | **z** | **p(uncorrected)** | **p(corrected)** | **Significance** |
| **Number of incorrect trials** | 0 | 0.084 | 0.096 | NA | 0.881 | 0.189 | 0.221 | n. s. |
|  | 6 | 0.173 | 0.113 | NA | 1.530 | 0.063 | 0.110 | n. s. |
|  | 12 | 0.192 | 0.108 | NA | 1.788 | 0.037 | 0.094 | * |
|  | 18 | 0.293 | 0.109 | NA | 2.687 | 0.004 | 0.025 | ** |
|  | 24 | -0.005 | 0.110 | NA | -0.048 | 0.519 | 0.519 | n. s. |
|  | 30 | 0.183 | 0.105 | NA | 1.746 | 0.040 | 0.094 | * |
|  | 36 | 0.151 | 0.122 | NA | 1.237 | 0.108 | 0.151 | n. s. |

MT: movement time; IPI: inter-press interval; ‘**’ significant FDR-corrected for number of sessions, q< 0.05; ‘*’ significant uncorrected, p < 0.05; ‘n. s.’ not significant.

**Supplementary Table 6. Statistics for cross-nobis dissimilarities of activation patterns.** Group (intervention/control) x practice (trained - untrained vs. untrained - untrained) interactions in cross-nobis dissimilarities between activation patterns of different sequences, for each MRI session and ROI (right hemisphere ROIs are shown Figure 4 A). A positive effect denotes a larger dissimilarity between neural patterns of trained and untrained sequences in the intervention group relative to the dissimilarity between neural patterns of pairs of untrained sequences and controls (cf. Figure 4B).

| **Label** | **MRI session** | **Estimate** | **Std. Error** | **df** | **t** | **p(unc.)** | **p(cor.)** | **p(cor. ROI)** | **Sign.** |
| --- | --- | --- | --- | --- | --- | --- | --- | --- | --- |
| **Left PM** | 0 | -0.022 | 0.012 | 31 | -1.827 | 0.966 | 0.984 | 0.966 | n. s. |
| **Left PM** | 1 | 0.009 | 0.015 | 48 | 0.587 | 0.279 | 0.390 | 0.325 | n. s. |
| **Left PM** | 2 | 0.044 | 0.012 | 52 | 3.562 | 1.84e-04 | 0.002 | 6.44e-04 | *** |
| **Left PM** | 3 | 0.037 | 0.014 | 54 | 2.594 | 0.005 | 0.018 | 0.008 | *** |
| **Left PM** | 4 | 0.034 | 0.010 | 54 | 3.365 | 3.83e-04 | 0.004 | 8.93e-04 | *** |
| **Left PM** | 5 | 0.049 | 0.013 | 51 | 3.709 | 1.04e-04 | 0.002 | 6.44e-04 | *** |
| **Left PM** | 6 | 0.030 | 0.015 | 42 | 2.008 | 0.022 | 0.057 | 0.031 | ** |
| **Left PS** | 0 | -0.021 | 0.012 | 31 | -1.699 | 0.955 | 0.984 | 0.955 | n. s. |
| **Left PS** | 1 | -0.005 | 0.009 | 48 | -0.520 | 0.698 | 0.800 | 0.815 | n. s. |
| **Left PS** | 2 | 0.002 | 0.010 | 52 | 0.259 | 0.398 | 0.518 | 0.696 | n. s. |
| **Left PS** | 3 | -3.73e-04 | 0.013 | 54 | -0.029 | 0.512 | 0.637 | 0.716 | n. s. |
| **Left PS** | 4 | 0.014 | 0.010 | 54 | 1.414 | 0.079 | 0.130 | 0.551 | n. s. |
| **Left PS** | 5 | 0.008 | 0.010 | 51 | 0.808 | 0.210 | 0.301 | 0.696 | n. s. |
| **Left PS** | 6 | 0.004 | 0.011 | 42 | 0.378 | 0.353 | 0.470 | 0.696 | n. s. |
| **Left SMA** | 0 | -0.024 | 0.015 | 31 | -1.600 | 0.945 | 0.984 | 0.945 | n. s. |
| **Left SMA** | 1 | 0.029 | 0.014 | 48 | 2.102 | 0.018 | 0.050 | 0.031 | *** |
| **Left SMA** | 2 | 0.023 | 0.012 | 52 | 1.850 | 0.032 | 0.069 | 0.045 | ** |
| **Left SMA** | 3 | 0.014 | 0.016 | 54 | 0.846 | 0.199 | 0.293 | 0.232 | n. s. |
| **Left SMA** | 4 | 0.033 | 0.013 | 54 | 2.568 | 0.005 | 0.018 | 0.018 | *** |
| **Left SMA** | 5 | 0.032 | 0.010 | 51 | 3.224 | 6.33e-04 | 0.004 | 0.004 | *** |
| **Left SMA** | 6 | 0.033 | 0.015 | 42 | 2.274 | 0.011 | 0.034 | 0.027 | *** |
| **Left SPL** | 0 | -0.021 | 0.012 | 31 | -1.780 | 0.962 | 0.984 | 0.962 | n. s. |
| **Left SPL** | 1 | 0.027 | 0.014 | 48 | 1.917 | 0.028 | 0.064 | 0.032 | ** |
| **Left SPL** | 2 | 0.028 | 0.012 | 52 | 2.423 | 0.008 | 0.025 | 0.012 | *** |
| **Left SPL** | 3 | 0.036 | 0.013 | 54 | 2.795 | 0.003 | 0.011 | 0.006 | *** |
| **Left SPL** | 4 | 0.035 | 0.011 | 54 | 3.065 | 0.001 | 0.006 | 0.004 | *** |
| **Left SPL** | 5 | 0.039 | 0.012 | 51 | 3.290 | 5.02e-04 | 0.004 | 0.004 | *** |
| **Left SPL** | 6 | 0.029 | 0.012 | 42 | 2.371 | 0.009 | 0.028 | 0.012 | *** |
| **Right PM** | 0 | -0.015 | 0.013 | 31 | -1.170 | 0.879 | 0.984 | 0.879 | n. s. |
| **Right PM** | 1 | 0.027 | 0.016 | 48 | 1.722 | 0.042 | 0.081 | 0.050 | ** |
| **Right PM** | 2 | 0.058 | 0.014 | 52 | 4.130 | 1.81e-05 | 5.08e-04 | 6.35e-05 | *** |
| **Right PM** | 3 | 0.050 | 0.016 | 54 | 3.071 | 0.001 | 0.006 | 0.002 | *** |
| **Right PM** | 4 | 0.041 | 0.013 | 54 | 3.220 | 6.42e-04 | 0.004 | 0.001 | *** |
| **Right PM** | 5 | 0.055 | 0.012 | 51 | 4.664 | 1.55e-06 | 8.68e-05 | 1.08e-05 | *** |
| **Right PM** | 6 | 0.037 | 0.019 | 42 | 1.977 | 0.024 | 0.058 | 0.034 | ** |
| **Right PS** | 0 | -0.040 | 0.018 | 31 | -2.150 | 0.984 | 0.984 | 0.984 | n. s. |
| **Right PS** | 1 | 0.018 | 0.018 | 48 | 1.000 | 0.159 | 0.240 | 0.555 | n. s. |
| **Right PS** | 2 | -0.007 | 0.014 | 52 | -0.525 | 0.700 | 0.800 | 0.817 | n. s. |
| **Right PS** | 3 | -0.002 | 0.017 | 54 | -0.108 | 0.543 | 0.661 | 0.817 | n. s. |
| **Right PS** | 4 | 0.015 | 0.015 | 54 | 1.015 | 0.155 | 0.240 | 0.555 | n. s. |
| **Right PS** | 5 | -0.004 | 0.012 | 51 | -0.309 | 0.621 | 0.740 | 0.817 | n. s. |
| **Right PS** | 6 | 0.009 | 0.019 | 42 | 0.454 | 0.325 | 0.444 | 0.758 | n. s. |
| **Right SMA** | 0 | -0.020 | 0.015 | 31 | -1.315 | 0.906 | 0.984 | 0.906 | n. s. |
| **Right SMA** | 1 | 0.030 | 0.017 | 48 | 1.747 | 0.040 | 0.081 | 0.082 | * |
| **Right SMA** | 2 | 0.035 | 0.012 | 52 | 2.973 | 0.001 | 0.007 | 0.010 | *** |
| **Right SMA** | 3 | 0.020 | 0.014 | 54 | 1.421 | 0.078 | 0.130 | 0.092 | n. s. |
| **Right SMA** | 4 | 0.020 | 0.014 | 54 | 1.410 | 0.079 | 0.130 | 0.092 | n. s. |
| **Right SMA** | 5 | 0.017 | 0.010 | 51 | 1.677 | 0.047 | 0.084 | 0.082 | * |
| **Right SMA** | 6 | 0.028 | 0.015 | 42 | 1.786 | 0.037 | 0.077 | 0.082 | * |
| **Right SPL** | 0 | 1.95e-04 | 0.013 | 31 | 0.016 | 0.494 | 0.628 | 0.494 | n. s. |
| **Right SPL** | 1 | 0.019 | 0.018 | 48 | 1.087 | 0.139 | 0.222 | 0.162 | n. s. |
| **Right SPL** | 2 | 0.031 | 0.015 | 52 | 2.084 | 0.019 | 0.050 | 0.043 | *** |
| **Right SPL** | 3 | 0.023 | 0.013 | 54 | 1.715 | 0.043 | 0.081 | 0.060 | * |
| **Right SPL** | 4 | 0.036 | 0.013 | 54 | 2.757 | 0.003 | 0.012 | 0.010 | *** |
| **Right SPL** | 5 | 0.023 | 0.012 | 51 | 1.893 | 0.029 | 0.065 | 0.051 | * |
| **Right SPL** | 6 | 0.048 | 0.013 | 42 | 3.637 | 1.38e-04 | 0.002 | 9.66e-04 | *** |

MRI session 0 refers to the baseline session; p(unc.): p-value, uncorrected; p(cor.): p-value, FDR-corrected corrected for number of sessions (q-value); p(cor. ROI): p-value, corrected for sessions and ROIs (q-value); Sign.: significance; ‘***’ significant FDR-corrected for number of sessions and ROIs, ‘**’ significant FDR-corrected for number of sessions, q< 0.05; ‘*’ significant uncorrected, p < 0.05; ‘n. s.’ not significant.

**Supplementary Table 7. Statistics for the variability index of the multivariate activation patterns.**

Group (intervention/control) x practice (trained/untrained) interactions in the index of variability of activation patterns, for each MRI session and ROI (right hemisphere ROIs are shown Figure 4 A). None of the interactions was significant after correcting for number of sessions and ROIs (cf. Supplementary Figure 5).

| **Label** | **MRI session** | **Estimate** | **Std. Error** | **df** | **t** | **p(unc.)** | **p(cor.)** | **p(cor. ROI)** | **Sign.** |
| --- | --- | --- | --- | --- | --- | --- | --- | --- | --- |
| **Left PM** | 0 | 0.009 | 0.029 | 31 | 0.303 | 0.619 | 0.737 | 0.619 | n. s. |
| **Left PM** | 1 | 0.005 | 0.023 | 48 | 0.204 | 0.581 | 0.707 | 0.619 | n. s. |
| **Left PM** | 2 | 0.002 | 0.021 | 52 | 0.106 | 0.542 | 0.707 | 0.619 | n. s. |
| **Left PM** | 3 | -0.050 | 0.037 | 54 | -1.354 | 0.088 | 0.400 | 0.224 | n. s. |
| **Left PM** | 4 | -0.012 | 0.022 | 54 | -0.554 | 0.290 | 0.676 | 0.507 | n. s. |
| **Left PM** | 5 | -0.045 | 0.034 | 51 | -1.305 | 0.096 | 0.400 | 0.224 | n. s. |
| **Left PM** | 6 | -0.060 | 0.026 | 42 | -2.311 | 0.010 | 0.117 | 0.073 | * |
| **Left PS** | 0 | -4.51e-05 | 0.032 | 31 | -0.001 | 0.499 | 0.707 | 0.598 | n. s. |
| **Left PS** | 1 | 7.85e-04 | 0.025 | 48 | 0.031 | 0.512 | 0.707 | 0.598 | n. s. |
| **Left PS** | 2 | 0.021 | 0.024 | 52 | 0.870 | 0.808 | 0.808 | 0.808 | n. s. |
| **Left PS** | 3 | -0.057 | 0.041 | 54 | -1.377 | 0.084 | 0.400 | 0.295 | n. s. |
| **Left PS** | 4 | -0.007 | 0.021 | 54 | -0.334 | 0.369 | 0.707 | 0.598 | n. s. |
| **Left PS** | 5 | -0.007 | 0.036 | 51 | -0.193 | 0.423 | 0.707 | 0.598 | n. s. |
| **Left PS** | 6 | -0.054 | 0.029 | 42 | -1.842 | 0.033 | 0.229 | 0.229 | * |
| **Left SMA** | 0 | 0.015 | 0.030 | 31 | 0.504 | 0.693 | 0.746 | 0.693 | n. s. |
| **Left SMA** | 1 | -1.57e-04 | 0.027 | 48 | -0.006 | 0.498 | 0.707 | 0.693 | n. s. |
| **Left SMA** | 2 | 0.011 | 0.026 | 52 | 0.440 | 0.670 | 0.737 | 0.693 | n. s. |
| **Left SMA** | 3 | -0.059 | 0.035 | 54 | -1.675 | 0.047 | 0.292 | 0.164 | * |
| **Left SMA** | 4 | -0.010 | 0.025 | 54 | -0.402 | 0.344 | 0.707 | 0.602 | n. s. |
| **Left SMA** | 5 | -0.023 | 0.040 | 51 | -0.569 | 0.285 | 0.676 | 0.602 | n. s. |
| **Left SMA** | 6 | -0.067 | 0.029 | 41.999 | -2.318 | 0.010 | 0.117 | 0.072 | * |
| **Left SPL** | 0 | 0.022 | 0.031 | 31 | 0.726 | 0.766 | 0.795 | 0.793 | n. s. |
| **Left SPL** | 1 | 0.019 | 0.023 | 48 | 0.816 | 0.793 | 0.807 | 0.793 | n. s. |
| **Left SPL** | 2 | 0.010 | 0.022 | 52 | 0.444 | 0.672 | 0.737 | 0.793 | n. s. |
| **Left SPL** | 3 | -0.037 | 0.031 | 54 | -1.206 | 0.114 | 0.425 | 0.349 | n. s. |
| **Left SPL** | 4 | -0.013 | 0.019 | 54 | -0.661 | 0.254 | 0.676 | 0.445 | n. s. |
| **Left SPL** | 5 | -0.032 | 0.031 | 51 | -1.039 | 0.149 | 0.492 | 0.349 | n. s. |
| **Left SPL** | 6 | -0.076 | 0.025 | 42 | -3.033 | 0.001 | 0.068 | 0.008 | ** |
| **Right PM** | 0 | 0.005 | 0.032 | 31 | 0.173 | 0.569 | 0.707 | 0.569 | n. s. |
| **Right PM** | 1 | 0.001 | 0.024 | 48 | 0.054 | 0.521 | 0.707 | 0.569 | n. s. |
| **Right PM** | 2 | -0.002 | 0.021 | 52 | -0.117 | 0.454 | 0.707 | 0.569 | n. s. |
| **Right PM** | 3 | -0.050 | 0.039 | 54 | -1.282 | 0.100 | 0.400 | 0.350 | n. s. |
| **Right PM** | 4 | -0.008 | 0.022 | 54 | -0.373 | 0.355 | 0.707 | 0.569 | n. s. |
| **Right PM** | 5 | -0.021 | 0.033 | 51 | -0.629 | 0.265 | 0.676 | 0.569 | n. s. |
| **Right PM** | 6 | -0.050 | 0.026 | 42 | -1.917 | 0.028 | 0.226 | 0.193 | * |
| **Right PS** | 0 | 0.018 | 0.029 | 31 | 0.627 | 0.735 | 0.776 | 0.735 | n. s. |
| **Right PS** | 1 | 0.005 | 0.024 | 48 | 0.197 | 0.578 | 0.707 | 0.674 | n. s. |
| **Right PS** | 2 | 0.004 | 0.023 | 52 | 0.192 | 0.576 | 0.707 | 0.674 | n. s. |
| **Right PS** | 3 | -0.028 | 0.036 | 54 | -0.778 | 0.218 | 0.643 | 0.674 | n. s. |
| **Right PS** | 4 | 0.003 | 0.023 | 54 | 0.152 | 0.560 | 0.707 | 0.674 | n. s. |
| **Right PS** | 5 | -0.012 | 0.035 | 51 | -0.345 | 0.365 | 0.707 | 0.674 | n. s. |
| **Right PS** | 6 | -0.062 | 0.025 | 42 | -2.453 | 0.007 | 0.117 | 0.050 | ** |
| **Right SMA** | 0 | 0.005 | 0.032 | 31 | 0.159 | 0.563 | 0.707 | 0.563 | n. s. |
| **Right SMA** | 1 | -0.012 | 0.024 | 48 | -0.503 | 0.307 | 0.688 | 0.538 | n. s. |
| **Right SMA** | 2 | 0.002 | 0.027 | 52 | 0.085 | 0.534 | 0.707 | 0.563 | n. s. |
| **Right SMA** | 3 | -0.053 | 0.038 | 54 | -1.404 | 0.080 | 0.400 | 0.281 | n. s. |
| **Right SMA** | 4 | -0.004 | 0.024 | 54 | -0.158 | 0.437 | 0.707 | 0.563 | n. s. |
| **Right SMA** | 5 | -0.023 | 0.039 | 51 | -0.585 | 0.279 | 0.676 | 0.538 | n. s. |
| **Right SMA** | 6 | -0.057 | 0.030 | 42 | -1.906 | 0.028 | 0.226 | 0.198 | * |
| **Right SPL** | 0 | 0.010 | 0.026 | 31 | 0.404 | 0.657 | 0.737 | 0.657 | n. s. |
| **Right SPL** | 1 | -7.76e-04 | 0.024 | 48 | -0.032 | 0.487 | 0.707 | 0.657 | n. s. |
| **Right SPL** | 2 | 0.008 | 0.020 | 52 | 0.393 | 0.653 | 0.737 | 0.657 | n. s. |
| **Right SPL** | 3 | -0.035 | 0.032 | 54 | -1.072 | 0.142 | 0.492 | 0.480 | n. s. |
| **Right SPL** | 4 | 0.002 | 0.019 | 54 | 0.118 | 0.547 | 0.707 | 0.657 | n. s. |
| **Right SPL** | 5 | -0.029 | 0.035 | 51 | -0.822 | 0.206 | 0.640 | 0.480 | n. s. |
| **Right SPL** | 6 | -0.063 | 0.027 | 42 | -2.331 | 0.010 | 0.117 | 0.069 | * |

MRI session 0 refers to the baseline session; p(unc.): p-value, uncorrected; p(cor.): p-value, FDR-corrected corrected for number of sessions (q-value); p(cor. ROI): p-value, corrected for sessions and ROIs (q-value); Sign.: significance; ‘***’ significant FDR-corrected for number of sessions and ROIs, ‘**’ significant FDR-corrected for number of sessions, q< 0.05; ‘*’ significant uncorrected, p < 0.05; ‘n. s.’ not significant.

**Supplementary Table 8. Statistics for the accuracy of a linear classifier of the multivariate activity patterns.** Group (intervention/control) x practice (trained - untrained vs. untrained - untrained) interactions in classification accuracy of activation patterns of different sequences, for each MRI session and ROI (right hemisphere ROIs are shown Figure 4 A). A positive effect denotes a larger accuracy when classifying between neural patterns of trained and untrained sequences in the intervention group relative to classifying neural patterns of pairs of untrained sequences and relative to controls (cf. Supplementary Figure 6).

| **Label** | **MRI session** | **Estimate** | **Std. Error** | **df** | **t** | **p(unc.)** | **p(cor.)** | **p(cor. ROI)** | **Sign.** |
| --- | --- | --- | --- | --- | --- | --- | --- | --- | --- |
| **Left PM** | 0 | -0.035 | 0.022 | 31 | -1.583 | 0.943 | 0.943 | 0.943 | n. s. |
| **Left PM** | 1 | 0.024 | 0.023 | 48 | 1.057 | 0.145 | 0.270 | 0.169 | n. s. |
| **Left PM** | 2 | 0.079 | 0.018 | 52 | 4.346 | 6.94e-06 | 3.88e-04 | 4.86e-05 | *** |
| **Left PM** | 3 | 0.067 | 0.019 | 54 | 3.523 | 2.14e-04 | 0.004 | 7.48e-04 | *** |
| **Left PM** | 4 | 0.058 | 0.022 | 54 | 2.683 | 0.004 | 0.019 | 0.009 | *** |
| **Left PM** | 5 | 0.054 | 0.022 | 51 | 2.489 | 0.006 | 0.026 | 0.011 | *** |
| **Left PM** | 6 | 0.058 | 0.024 | 42 | 2.392 | 0.008 | 0.029 | 0.012 | *** |
| **Left PS** | 0 | 0.006 | 0.020 | 31 | 0.305 | 0.380 | 0.496 | 0.650 | n. s. |
| **Left PS** | 1 | 0.010 | 0.017 | 48 | 0.563 | 0.287 | 0.401 | 0.650 | n. s. |
| **Left PS** | 2 | -0.020 | 0.016 | 52 | -1.183 | 0.882 | 0.910 | 0.882 | n. s. |
| **Left PS** | 3 | 0.002 | 0.018 | 54 | 0.090 | 0.464 | 0.585 | 0.650 | n. s. |
| **Left PS** | 4 | 0.020 | 0.018 | 54 | 1.149 | 0.125 | 0.251 | 0.650 | n. s. |
| **Left PS** | 5 | -0.017 | 0.021 | 51 | -0.801 | 0.788 | 0.883 | 0.882 | n. s. |
| **Left PS** | 6 | 0.014 | 0.021 | 42 | 0.657 | 0.256 | 0.377 | 0.650 | n. s. |
| **Left SMA** | 0 | -0.018 | 0.019 | 31 | -0.980 | 0.837 | 0.901 | 0.837 | n. s. |
| **Left SMA** | 1 | 0.014 | 0.018 | 48 | 0.810 | 0.209 | 0.334 | 0.244 | n. s. |
| **Left SMA** | 2 | 0.054 | 0.021 | 52 | 2.562 | 0.005 | 0.022 | 0.018 | *** |
| **Left SMA** | 3 | 0.018 | 0.018 | 54 | 1.019 | 0.154 | 0.270 | 0.216 | n. s. |
| **Left SMA** | 4 | 0.037 | 0.019 | 54 | 1.969 | 0.024 | 0.061 | 0.043 | ** |
| **Left SMA** | 5 | 0.057 | 0.018 | 51 | 3.215 | 6.52e-04 | 0.006 | 0.005 | *** |
| **Left SMA** | 6 | 0.042 | 0.020 | 42 | 2.060 | 0.020 | 0.058 | 0.043 | ** |
| **Left SPL** | 0 | -0.034 | 0.030 | 31 | -1.137 | 0.872 | 0.910 | 0.872 | n. s. |
| **Left SPL** | 1 | 0.046 | 0.023 | 48 | 1.991 | 0.023 | 0.061 | 0.041 | ** |
| **Left SPL** | 2 | 0.015 | 0.021 | 52 | 0.719 | 0.236 | 0.357 | 0.275 | n. s. |
| **Left SPL** | 3 | 0.045 | 0.021 | 54 | 2.215 | 0.013 | 0.044 | 0.031 | *** |
| **Left SPL** | 4 | 0.046 | 0.019 | 54 | 2.443 | 0.007 | 0.027 | 0.025 | *** |
| **Left SPL** | 5 | 0.032 | 0.020 | 51 | 1.575 | 0.058 | 0.129 | 0.081 | n. s. |
| **Left SPL** | 6 | 0.073 | 0.022 | 42 | 3.277 | 5.25e-04 | 0.006 | 0.004 | *** |
| **Right PM** | 0 | -0.013 | 0.022 | 31 | -0.583 | 0.720 | 0.840 | 0.720 | n. s. |
| **Right PM** | 1 | 0.007 | 0.022 | 48 | 0.302 | 0.381 | 0.496 | 0.445 | n. s. |
| **Right PM** | 2 | 0.060 | 0.019 | 52 | 3.117 | 9.14e-04 | 0.006 | 0.003 | *** |
| **Right PM** | 3 | 0.045 | 0.023 | 54 | 1.980 | 0.024 | 0.061 | 0.033 | ** |
| **Right PM** | 4 | 0.060 | 0.023 | 54 | 2.575 | 0.005 | 0.022 | 0.009 | *** |
| **Right PM** | 5 | 0.061 | 0.018 | 51 | 3.335 | 4.26e-04 | 0.006 | 0.003 | *** |
| **Right PM** | 6 | 0.053 | 0.020 | 42 | 2.696 | 0.004 | 0.019 | 0.008 | *** |
| **Right PS** | 0 | -0.018 | 0.028 | 31 | -0.634 | 0.737 | 0.842 | 0.828 | n. s. |
| **Right PS** | 1 | 0.032 | 0.024 | 48 | 1.356 | 0.088 | 0.189 | 0.613 | n. s. |
| **Right PS** | 2 | -0.019 | 0.020 | 52 | -0.946 | 0.828 | 0.901 | 0.828 | n. s. |
| **Right PS** | 3 | 0.012 | 0.019 | 54 | 0.611 | 0.271 | 0.388 | 0.631 | n. s. |
| **Right PS** | 4 | 0.017 | 0.019 | 54 | 0.878 | 0.190 | 0.322 | 0.631 | n. s. |
| **Right PS** | 5 | 0.002 | 0.023 | 51 | 0.075 | 0.470 | 0.585 | 0.731 | n. s. |
| **Right PS** | 6 | -0.001 | 0.026 | 42 | -0.055 | 0.522 | 0.635 | 0.731 | n. s. |
| **Right SMA** | 0 | -0.029 | 0.023 | 31 | -1.247 | 0.894 | 0.910 | 0.894 | n. s. |
| **Right SMA** | 1 | 0.020 | 0.017 | 48 | 1.173 | 0.120 | 0.250 | 0.305 | n. s. |
| **Right SMA** | 2 | 0.058 | 0.018 | 52 | 3.134 | 8.61e-04 | 0.006 | 0.006 | *** |
| **Right SMA** | 3 | 0.016 | 0.021 | 54 | 0.780 | 0.218 | 0.338 | 0.305 | n. s. |
| **Right SMA** | 4 | 0.010 | 0.021 | 54 | 0.453 | 0.325 | 0.444 | 0.380 | n. s. |
| **Right SMA** | 5 | 0.018 | 0.017 | 51 | 1.034 | 0.151 | 0.270 | 0.305 | n. s. |
| **Right SMA** | 6 | 0.016 | 0.019 | 42 | 0.818 | 0.207 | 0.334 | 0.305 | n. s. |
| **Right SPL** | 0 | -0.013 | 0.025 | 31 | -0.508 | 0.694 | 0.827 | 0.694 | n. s. |
| **Right SPL** | 1 | 0.062 | 0.022 | 48 | 2.850 | 0.002 | 0.014 | 0.008 | *** |
| **Right SPL** | 2 | 0.050 | 0.023 | 52 | 2.195 | 0.014 | 0.044 | 0.033 | *** |
| **Right SPL** | 3 | 0.024 | 0.022 | 54 | 1.062 | 0.144 | 0.270 | 0.168 | n. s. |
| **Right SPL** | 4 | 0.043 | 0.022 | 54 | 1.960 | 0.025 | 0.061 | 0.044 | ** |
| **Right SPL** | 5 | 0.037 | 0.021 | 51 | 1.790 | 0.037 | 0.086 | 0.051 | * |
| **Right SPL** | 6 | 0.074 | 0.021 | 42 | 3.494 | 2.38e-04 | 0.004 | 0.002 | *** |

MRI session 0 refers to the baseline session; p(unc.): p-value, uncorrected; p(cor.): p-value, FDR-corrected corrected for number of sessions (q-value); p(cor. ROI): p-value, corrected for sessions and ROIs (q-value); Sign.: significance; ‘***’ significant FDR-corrected for number of sessions and ROIs, ‘**’ significant FDR-corrected for number of sessions, q< 0.05; ‘*’ significant uncorrected, p < 0.05; ‘n. s.’ not significant.
